# Comparative Population Genomics of Bread Wheat *(Triticum aestivum)* Reveals Its Cultivation and Breeding History in China

**DOI:** 10.1101/519587

**Authors:** Haofeng Chen, Chengzhi Jiao, Ying Wang, Yuange Wang, Caihuan Tian, Haopeng Yu, Jing Wang, Xiangfeng Wang, Fei Lu, Xiangdong Fu, Yongbiao Xue, Wenkai Jiang, Hongqing Ling, Hongfeng Lu, Yuling Jiao

## Abstract

The evolution of bread wheat *(Triticum aestivum)* is distinctive in that domestication, natural hybridization, and allopolyploid speciation have all had significant effects on the diversification of its genome. Wheat was spread around the world by humans and has been cultivated in China for ~4,600 years. Here, we report a comprehensive assessment of the evolution of wheat based on the genome-wide resequencing of 120 representative landraces and elite wheat accessions from China and other representative regions. We found substantially higher genetic diversity in the A and B subgenomes than in the D subgenome. Notably, the A and B subgenomes of the modern Chinese elite cultivars were mainly derived from European landraces, while Chinese landraces had a greater contribution to their D subgenomes. The duplicated copies of homoeologous genes from the A, B, and D subgenomes were commonly found to be under different levels of selection. Our genome-wide assessment of the genetic changes associated with wheat breeding in China provides new strategies and practical targets for future breeding.

## Main

Bread wheat (*Triticum aestivum)* differs from other major grain crops, such as maize *(Zea mays)*, rice *(Oryza sativa)*, and barley *(Hordeum vulgare)*, by having an allohexaploid genome with six sets of chromosomes, two sets each from three closely related ancestral species that formed the A, B, and D subgenomes. Archaeological and genetic evidence indicated that wheat underwent two polyploidization events during its domestication. The first occurred ~0.82 million years ago between two diploid species, *T. urartu* (AA) and an unknown close relative of *Aegilops speltoides* (SS), which produced the allotetraploid *T. turgidum* (AABB). The second polyploidization occurred during cultivation around 8,000–10,000 years ago, when *T. turgidum* crossed with another diploid grass, *Ae. tauschii* (DD), to form the ancestral bread wheat *(T. aestivum*, AABBDD)^1–4^ In addition, a recent genome analysis suggested that *Ae. tauschii* originated from a more ancient homoploid hybridization event between the A and B lineage ancestors^5^. The A, B, and D subgenomes comprise the ~17-Gb allohexaploid bread wheat genome. Enormous genome sequencing efforts have resulted in the recent publication of a reference genome sequence for wheat^4,6–9^, which has greatly enhanced our understanding of the genome of this vital crop.

After its evolution in the Middle East and Mediterranean regions, bread wheat gradually spread to the rest of the world and was domesticated for human use^10^. China has been cultivating bread wheat for ~4,600 years^11,12^, and has been the largest wheat-producing country for more than two decades. Wheat has been under continuous artificial selection in the diverse ecological zones of China for thousands of years^11,13^; thus, the domestication and breeding of bread wheat in this country provides unique evolutionary insights into how its genome diversity was used and altered to meet the changing needs of the human population.

Recent advances in wheat genome sequencing have made pangenome studies possible^14,15^. To understand wheat genome diversity and evolution, we resequenced 120 landraces and elite bread wheat accessions at above a ten-fold coverage, and employed the resequencing data to unravel the genealogical history of wheat domestication. We identified the hybrid origins of various wheat cultivars and landraces, especially those cultivated in China. Using these data, we began to elucidate the genomic signatures that underlie human selection during the history of wheat breeding.

## Results

### Characterization of molecular diversity

We selected 120 wheat accessions for our study, including 95 landraces and 25 cultivars. The 60 Chinese accessions are representative ones of a Chinese mini core collection (Supplementary Fig. 1a and b), which covers the widest genome diversity of 23,090 Chinese landraces and cultivars^16^. The other accessions were selected based on genotyping-by-sequencing results of 326 representative species collected worldwide (Supplementary Fig. 1c). Notably, we selected 16 accessions from the regions of the Fertile Crescent where wheat was first cultivated. Thus, the resequenced accessions represent the widest genome diversity of both Chinese and worldwide wheat populations. The geographic distributions of these accessions included East Asia (China and Japan), West Asia, Central and South Asia, Europe, and America (Fig. 1a and Supplementary Table 1).

**Fig. 1.**
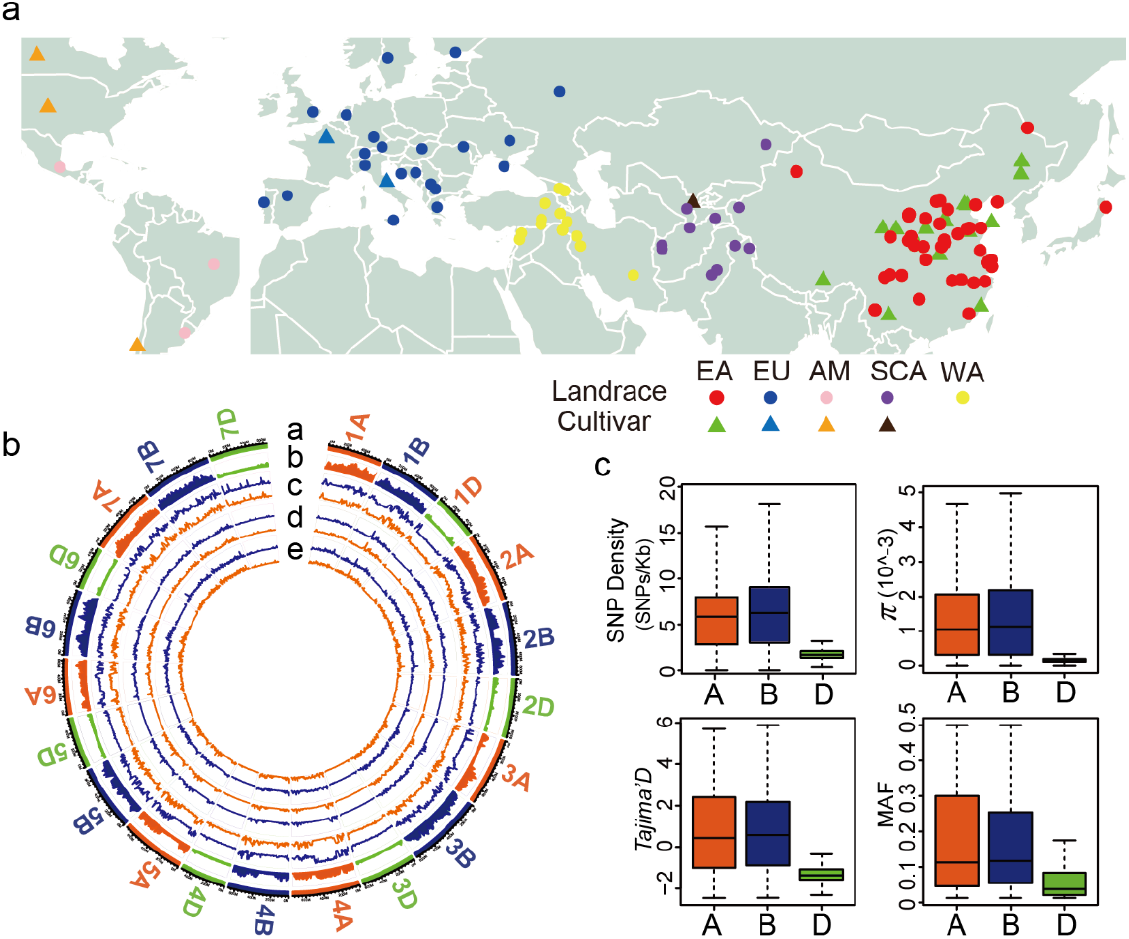
Characterization of SNP distribution across the A, B, and D subgenome chromosomes. **(a)** The geographic distribution of the 120 wheat accessions used in this study. The circles represent landraces, while triangles represent cultivars. Different colors represent populations from different geographic regions. EA, East Asia; WA, West Asia; EU, Europe; AM, America; SCA, South and Central Asia. **(b)** The genetic diversity of different populations along the chromosomes. From a to e, the sections respectively represent the chromosomes, SNP density, neutral evolutionary parameters (Tajima’s D), nucleotide diversity parameters (θ_π_), and Watterson’s θ, respectively. The red and blue lines in sections c to e represent the cultivars and landraces, respectively. **(c)** Comparison of the genomic characteristics of the A, B, and D subgenomes. MAF, minor allele frequencies.

A total of 21,676 Tb of sequence data was generated from the 120 accessions using Illumina paired-end short-read technology, with an average read depth of 11.04× for each individual (Supplementary Table 2). The reads were mapped against the IWGSC RefSeq v1.0 assembly of the wheat genome^9^ to identify genomic variants, yielding a ~90% coverage of the genome by at least four reads for each wheat accession. We detected a total of 70,172,600 single-nucleotide polymorphisms (SNPs) using the SAMtools software^17^, with an average density of 4.82 SNPs per kb. Most of these SNPs were located in intergenic regions (94.76%), with only 1.43% located in exonic regions (Table 1 and Supplementary Fig. 2). Of the SNPs located in exonic regions, 36.62% were synonymous and 54.80% were non-synonymous, with a non-synonymous/synonymous (N/S) ratio of 1.5. This N/S ratio is higher than was previously reported for other plants such as sorghum *(Sorghum bicolor*; N/S ratio = 1. 0)^18^, rice (N/S ratio = 1.2)^19^, soybean *(Glycine max*; N/S ratio = 1.47)^20^, and *Arabidopsis thaliana* (N/S ratio = 0.83)^21^.

**Table 1.** Summary of the SNP distribution in the A, B, and D subgenomes.

| Sub-genome | Up-stream<sup>a</sup> | Exonic |  |  |  |  |  | Intronic | 3' UTR | 5' UTR | 5' UTR; 3' UTR<sup>b</sup> | Splicing | Down-stream<sup>c</sup> | Up/down Stream<sup>d</sup> | Intergenic |
| --- | --- | --- | --- | --- | --- | --- | --- | --- | --- | --- | --- | --- | --- | --- | --- |
|  |  | Stop gain | Stop loss | Synonymous | Non-synonymous | N/S ratio | Unknowns |  |  |  |  |  |  |  |  |
| A | 325,784 | 8,180 | 1,782 | 133,267 | 207,728 | 1.56 | 27,325 | 366,052 | 11,108 | 12,388 | 1 | 1,466 | 302,952 | 46,595 | 27,276,394 |
| B | 349,547 | 9,130 | 1,953 | 141,373 | 224,779 | 1.59 | 30,931 | 394,269 | 11,557 | 14,315 | 11 | 1,615 | 324,819 | 52,059 | 31,395,654 |
| D | 124,077 | 3,487 | 644 | 82,817 | 105,693 | 1.28 | 8,690 | 151,873 | 5,590 | 5,881 | 4 | 660 | 117,057 | 18,823 | 7,287,737 |
| Unknown | 11,337 | 327 | 58 | 6,529 | 9,168 | 1.40 | 838 | 8,961 | 446 | 299 | 0 | 44 | 10,984 | 1,397 | 536,205 |
| Total | 810,745 | 21,124 | 4,437 | 363,986 | 547,368 | 1.50 | 67,784 | 921,155 | 28,701 | 32,883 | 16 | 3,785 | 755,812 | 118,874 | 66,495,990 |

<sup>a</sup>Regions within the 1 kb before the start codon of the downstream gene.
<sup>b</sup>Regions considered as both the 3'UTR of the upstream gene and 5'UTR of the downstream gene.
<sup>c</sup>Regions within the 1 kb after the stop codon of the upstream gene.
<sup>d</sup>Regions within the 1 kb after the stop codon of the upstream gene and also within the 1 kb before the start codon of the downstream gene.

We found that the A and B subgenomes harbor similar numbers of SNPs, with the B subgenome (~32.95 M, 6.36 SNPs per kb) containing slightly more than the A subgenome (~28.72 M, 5.82 SNPs per kb). By contrast, the D subgenome included only ~7.91 M SNPs and had the lowest SNP density (2.00 SNPs per kb) (Table 1, Fig. 1c). A previous finding showed that low numbers of polymorphic loci in the D subgenome may be a specific attribute of wheat, not of the D subgenome progenitor *Ae. tauschii^22^.* In all three wheat subgenomes, the majority of SNPs were located in intergenic regions, and non-synonymous SNPs were more common than synonymous SNPs (the N/S ratios of A, B, and D subgenomes were 1.56, 1.59, and 1.28, respectively; Table 1). In the wheat reference genome (IWGSC RefSeq Annotation v1.0), the gene models were classified into high-confidence (HC) and low-confidence (LC) genes based on their predicted sequence homology to gene products in the public database. In the coding regions, 47.78% of the SNPs were located in HC genes. These SNPs also had N/S ratios greater than 1 (Supplementary Table 3).

To measure the degree of polymorphism in the wheat genome, we measured the nucleotide diversity parameters π and Watterson’s *θ.* The overall wheat genomic nucleotide diversity (*π*) was 1.05 × 10^−3^, and Watterson’s *θ* was 0.84 × 10^−3^, which is consistent with a previous report (π = 0.8 × 10^−3^)^23^. The nucleotide diversity parameter for wheat was lower than for other major crops, including maize (π = 6.6 × 10^−3^)^24^, rice (π = 2.29 × 10^−3^)^25^, and sorghum (π = 2.4 × 10^−3^)^26^. The π value of the cultivars (0.92 × 10^−3^) was only slightly lower than that of the landraces (1.04 × 10^−3^). Similarly, the Watterson’s *θ* value for the cultivars (0.84 × 10^−3^) was slightly lower than that of the landraces (0.87 × 10^−3^). The mean fixation index value (*F*_ST_) between the cultivars and landraces was 0.03, suggesting limited population divergence between the cultivars and landraces.

We then separated the data into the A, B, and D subgenomes to understand their individual genetic diversities (Fig. 1b and Supplementary Table 4). In the individual A, B, and D subgenomes, the nucleotide diversity parameters π and Watterson’s *θ* were similar between the landraces and cultivars, indicating that the modern wheat cultivars retained most of the genetic diversity of the landraces during their domestication. The π and Watterson’s *θ* values each demonstrated that the A and B subgenomes had similar nucleotide diversities, while the D subgenome had only ~20% of the genetic diversity of the A or B subgenomes (Fig. 1c), consistent with a previous report^15^. The heterozygous rates of all accessions were low (average: 0.1655%), reflecting the lack of cross-pollination, consistent with the cleistogamy of wheat flowers. Among the three subgenomes, the D subgenome had a substantially lower heterozygous rate than the A and B subgenomes (Supplementary Table 5), consistent with its lower genetic diversity.

Tajima’s *D* was calculated to assess whether the observed nucleotide diversities showed evidence of deviation from neutrality. Some regions were significantly different from zero, which indicates that they may be under sweep selection (Fig. 1b). Of these, the *D* values from the A and B subgenomes were mostly positive, whereas those of the D subgenome were generally negative (Fig. 1c and Supplementary Table 4). Negative Tajima’s *D* values indicate the presence of low-frequency SNPs in the D subgenome, while the positive Tajima’s *D* values indicate a predominance of intermediate-frequency SNPs in the A and B subgenomes. The minor allele frequency (MAF) in the A, B, and D subgenomes significantly correlated with the Tajima’s *D* values (Supplementary Table 6). Because the three subgenomes were expected to have experienced a similar evolution history in the allohexaploid wheat after the second polyploidization event, the dramatic difference between D subgenome and the A and B subgenomes indicates an asymmetric selection history during domestication. Gene flow to bread wheat from wild and/or cultivated tetraploid *T. turgidum* (AABB genome) is likely common, as suggested by the identification of hybrid swarms between wild emmer wheat *(T. turgidum* subsp. *dicoccoides)* and bread wheat^27,28^. By contrast, the barrier to gene flow from *Ae. tauschii* (DD genome) to bread wheat is much more difficult to overcome. Nevertheless, this difference may also contribute to the variations in the evolution rates between the subgenomes.

### Population structure

To explore the phylogenetic relationships among the 120 wheat accessions, we constructed a phylogenetic tree using the neighbor-joining algorithm, based on the pairwise genetic distances between each accession determined using the SNP information. The phylogenetic analysis clustered the wheat accessions into two groups. Group 1 incorporated most of the European landraces, the West Asian landraces (marked as “Origin”), a few South and Central Asian landraces, and the majority of the East Asian cultivars (Fig. 2a). In group 2, most of the East Asian landraces were clustered together alongside some of the West Asian landraces and the majority of the South and Central Asian landraces (Fig. 2a). These two clearly defined groups were further strengthened by the results of a principal component analysis (PCA) and the Bayesian model-based clustering method (Fig. 2b). At *K* = 2, the group 1 accessions formed one cluster, while those of group 2 formed the other cluster. At the optimal presumed number of ancestral populations (K = 3), the landraces near the origin site (West, South, and Central Asian accessions) were separated from the main groups. At *K* = 4, the West Asian and South and Central Asian accessions were clearly divided. At *K* = 5, we obtained more refined clusters associated with the geographic distributions (Fig. 2c and Supplementary Fig. 5). These findings were consistent with the geographic documentation of wheat: from its domestication in the Fertile Crescent, bread wheat was transported west to Europe and America, and east to South and Central Asia and finally to East Asia via separate routes (Supplementary Fig. 4)^10^. Our phylogenetic analysis, PCA, and clustering approaches revealed that only a few Chinese cultivars are closely related to the Chinese landraces (Fig. 2a and B). The population structure revealed that the Chinese cultivars are clearly related to the European (and American) landraces when *K* = 2–5 (Fig. 2c). It is therefore evident that the Chinese landraces contributed a minor portion of the genetic diversity of the Chinese cultivars.

**Fig. 2.**
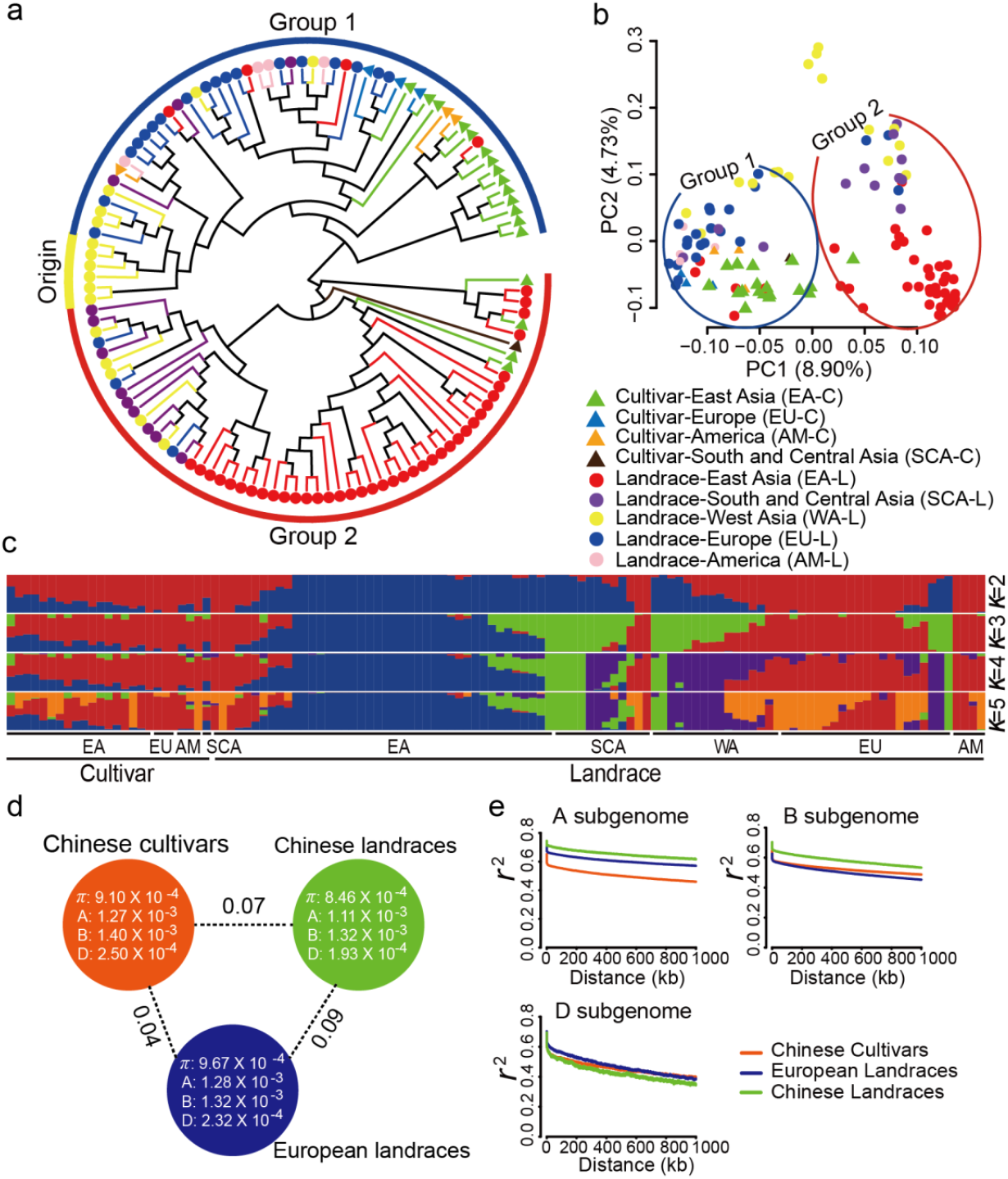
Population structure of the 120 wheat accessions. **(a)** Phylogenetic tree of the wheat accessions used in this study. The triangles represent wheat cultivars, while circles represent the landraces. The different colors represent populations from different geographic locations. “Origin” represents West Asia, which is considered the origin of bread wheat. “Group 1” represents the genotypes derived from the westward migration of bread wheat, including the European and American landraces, and the cultivars from Europe, America, and China. “Group 2” represents the groups derived from the eastward migration of bread wheat, including the South, Central, and East Asian landraces. **(b)** Principal component analysis of the 120 wheat accessions. **(a)** and **(b)** share the same key. **(c)** Population structure of the 120 wheat accessions. (d) Nucleotide diversity *(π)* and population divergence (*F*_ST_) across the Chinese cultivars, Chinese landraces, and European landraces. The values in each circle represent a measure of nucleotide diversity for this group and each of its subgenomes, while the values on each line indicate the population divergences between the two compared groups. **(e)** Decay of linkage disequilibrium (LD) in the A, B, and D subgenomes of three populations.

To understand whether different diploid ancestors contributed similar amounts of genetic diversity to the subgenomes, we further analyzed the population structure of each subgenome separately. The A and B subgenome population structures confirmed the evolutionary patterns observed for the whole genome (Supplementary Figs. 6 and 7). In contrast, the D subgenomes of most Chinese cultivars were closely related to the Chinese landraces in the phylogenetic tree analysis, which was further supported by the results of the PCA and genetic structure analysis (Supplementary Fig. 9). Taken together, the A and B subgenomes of the Chinese cultivars may mainly come from European landraces and cultivars with a genetic admixture of Chinese landraces, whereas the D subgenome contains more of a contribution from the Chinese landraces with a lesser admixture of European landraces.

To investigate the differences in genetic diversity between the three populations (Chinese cultivars, Chinese landraces, and European landraces), we calculated the nucleotide diversities *(π)* and found no significant difference among the three populations. The diversity of Chinese cultivars was slightly higher than that of the Chinese landraces, but slightly lower than that of the European landraces (Fig. 2d). We calculated the *F*_ST_ values to investigate the population divergences, which revealed that the divergence between the Chinese cultivars and European landraces (*F*_ST_ = 0.04) was smaller than the divergence of the Chinese cultivars and Chinese landraces (*F*_ST_ = 0.07). This analysis indicated a small population divergence (Fig. 2d).

We further analyzed the linkage disequilibrium (LD) of the three populations. The mean r^2^ between 0 and 1000 kb from the three populations was greater than 0.3, suggesting that wheat has a longer LD decay than cultivated rice (123 kb for *indica* varieties) and cultivated maize (30 kb)^29,30^. The mean r^2^ between 0 and 1000 kb of the D subgenome decreased more rapidly than for the A and B subgenomes, suggesting a lower level of selection on the D subgenome (Fig. 2e). The extent of LD in the A, B, and D subgenomes differed between the Chinese cultivars, Chinese landraces, and European landraces. For the A subgenome, the LD decay of the Chinese landraces was slightly higher than that of the European landraces, and significantly higher than that of the Chinese cultivars. For the B subgenome, the Chinese landraces also had the highest LD levels, with the Chinese cultivars and European landraces displaying similar levels of LD decay. In the D subgenome however, the LD decay was similar for all three populations. Different chromosomes had specific patterns of LD (Supplementary Fig. 3), which may be correlated with differences in heterochromatin levels^31^ and selection pressure.

### Introgressions from Chinese landraces and European varieties in the Chinese cultivars

We used a TreeMix analysis to infer ancient gene flows. A maximum-likelihood (ML) tree without migration events grouped the wheat accessions into seven clusters, which were similar to the above population structure patterns. Furthermore, we detected strong migration events in three clusters, namely the Chinese cultivars, Chinese landraces, and European cultivars. There were strong gene flows from the Chinese landraces to the Chinese cultivars, and from the European landraces to the European cultivars (Supplementary Fig. 6a and b). We next individually analyzed the three subgenomes. The ML trees for the A and B subgenomes of the accessions had similar topological structures to the ML tree using the whole genomes (Fig. 3a). We also found strong gene flows from the Chinese landraces to the Chinese cultivars and from the ancient western landraces to the European landraces in both the A and B subgenomes (Fig. 3a). When the D subgenome was considered, the accessions formed a different ML tree structure, with Chinese landraces grouped with Chinese cultivars, and clear gene flows from the European cultivars to the Chinese cultivars, and from the South and Central Asian landraces to the European landraces (Fig. 3a).

**Fig. 3.**
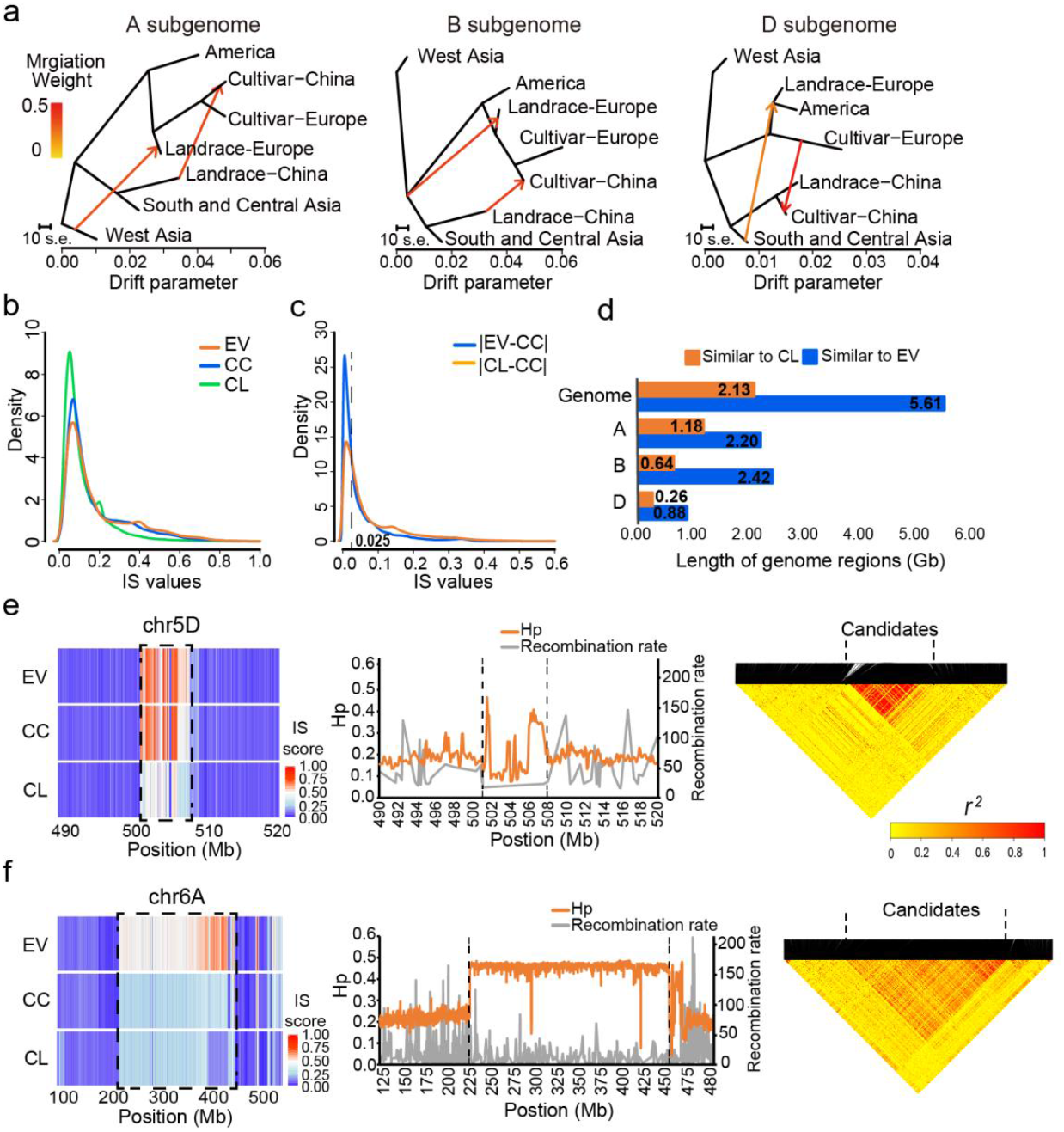
Haplotype introgressions from the European and Chinese landraces into the Chinese cultivars. **(a)** Inferred phylogenetic trees of the A, B, and D subgenomes of different populations, including mixture events. Horizontal branch lengths are proportional to the amount of genetic drift. Migration arrows are colored according to their weight. **(b)** Density distribution of identity score (IS) values for three populations. **(c)** Density distribution of the difference in IS values between two populations. lEL-CC| represents the absolute value of the IS differences between the European varieties and the Chinese cultivars. |CL-CC| represents the absolute value of the IS differences between the Chinese landraces and the Chinese cultivars. **(d)** Amount of similar genomic regions between populations. Orange represents the length of similar genomic regions between the Chinese cultivars and the Chinese landraces, and blue represents the length of similar genomic regions between the Chinese cultivars and the European varieties. The statistical results of the entire genome (Genome) as well as the individual A, B, and D subgenomes. The numbers represent the total length of the similar genomic regions (Gb). **(e)** Candidate introgression region between the European varieties and the Chinese cultivars. The left panel presents details of the 8-Mb candidate region, located on chromosome 5D, in which the Chinese cultivars are more similar to the European varieties than the Chinese landraces. The middle panel represents the distribution of the population heterozygosity (H_P_) and recombination rates in this candidate region. The right panel represents the LD heatmap of this candidate region. **(f)** Candidate introgression region (200 Mb, located on chromosome 6A) between the Chinese landraces and Chinese cultivars. The central and right panels are as described for (e).

To further elucidate the contributions from the European varieties and Chinese landraces to the Chinese cultivars, we used an identity score (IS) analysis^32^ to scan each chromosome. The IS analyses for the three groups (European varieties, Chinese landraces, and Chinese cultivars) revealed that the Chinese landraces had more similarity to the reference genome (Chinese Spring, a Chinese landrace) than the European and Chinese cultivars, while the Chinese cultivars were more similar to the European varieties than the Chinese landraces (Fig. 3b). Consistent with the results of the population structure analysis, the IS heatmap showed that the European varieties contributed more to the Chinese cultivars than did the Chinese landraces (Supplementary Fig. 10). More differences were detected between the Chinese landraces and the Chinese cultivars than between the European varieties and the Chinese cultivars when using IS values less than 0.0025 as the threshold (Fig. 3c). Specifically, 280,299 regions were detected, spanning 5.61 Gb (39.28% of the genomes), in which the Chinese cultivars and European varieties were more similar than the Chinese cultivars and Chinese landraces, while only 106,453 regions, spanning 2.13 Gb (14.92% of the genomes) were identified in which Chinese cultivars and Chinese landraces were more similar (Fig. 3d). Within most chromosomes, we observed similar patterns of homology, although there were clear variations (Supplementary Fig. 11, Supplementary Table 7a).

We were particularly interested in the many exceptionally large dispersed introgression regions identified in the genomes of the wheat population. Between the Chinese landraces and the Chinese cultivars, the introgression regions are mainly located in the A and B subgenomes, while those between the European varieties and the Chinese cultivars were mainly located in the D subgenome. These introgression regions usually show higher heterozygosity, low recombination rates, and long-range LD^27^; for example, an 8-Mb region in chromosome 5D of the Chinese cultivars is enriched with SNPs from the European varieties (Fig. 3e). Within this region, the Chinese cultivars are highly heterozygous and have low recombination rates, as well as a significantly higher LD level (Fig. 3e). We focused on the mutation sites within this candidate introgression region, which covers 346 genes (Supplementary Table 7b). Using a Kyoto Encyclopedia of Genes and Genomes (KEGG) analysis, we observed a significant enrichment of genes involved in pathways related to photosynthesis within this region (Supplementary Table 7d), including genes involved in photosystem I *(TraesCS5D01G552400LC/PSAB, TraesCS5D01G552500LC/PSAB*, and *TraesCS5D01G552600LC/PSAB)* and photosystem II *(TraesCS5D01G550000LC/PSBA, TraesCS5D01G458100/PSBK, TraesCS5D01G458200/PSBI, TraesCS5D01G550600LC/PSBA, TraesCS5D01G550700LC/PSBC, TraesCS5D01G550800LC/PSBC, TraesCS5D01G458300/PSBZ*, and *TraesCS5D01G458400/PSBM).* Another 200-Mb region on chromosome 6A of the Chinese cultivars is enriched with SNPs from the Chinese landraces (Fig. 3f). This region also has higher heterozygosity and lower recombination rates within these lines, and shows a significantly higher LD (Fig. 3f). The 2,131 genes within this region (Supplementary Table 7c) were also enriched for oxidative phosphorylation and ribosome pathway functions (Supplementary Table 7e). Taken together, our IS analysis revealed the detailed contributions of the European landraces and Chinese landraces to the Chinese cultivars. Importantly, most of these regions are genome-specific, suggesting the functional diversification of the three genomes.

### Selective signals during diversification and modern breeding

To identify the genomic regions most affected by selection during wheat diversification and modern breeding in China, we scanned the patterns of genetic variation along the chromosomes on the basis of the 70.12 million SNPs. We calculated the nucleotide diversity ratio (θπ landrace/θπ cultivar) and genetic differentiation (*F*_ST_) between the cultivars and landraces. Long stretches of elevated *F*_ST_ were found along chromosome 4A, which is known to contain structurally rearranged chromosomal regions^33^. After excluding these rearranged regions, the top 5% of regions (~14.98 Mb) were considered putative sweep-selected sections, which contained 842 genes (Supplementary Table 8a). A comparison among the subgenomes indicated that selection in the D subgenome is substantially lower than in the A and B subgenomes, as indicated by the lower *F*_ST_ and nucleotide diversity ratio (θ_π landrace_/θ_π cultivar_) for the D subgenome (Fig. 1b). The A subgenome had the most selected regions (Supplementary Table 8b). We also used the XP-EHH^34^ and XP-CLR^35^ methods to identify the regions under sweep selection, which respectively led to the identification of 2.65 Mb (948 genes) and 16.63 Mb (2,168 genes) of the putatively selected regions (Supplementary Table 8c and d). The combination of the results from all approaches led to the identification of a total of 3,659 genes, the majority of which were located on the A and B subgenomes (Supplementary Table 8f). The selected regions contain many previously reported quantitative trait loci (QTL); for example, one on chromosome 6B and one on chromosome 7D both associated with the length of the uppermost internode length^36^. Gene Ontology (GO) and KEGG enrichment analyses revealed that the selected regions are enriched in genes with decarboxylating activity and glycine catabolism, suggesting that the carbon metabolic pathways were targeted during modern breeding. Additional pathways, such as selenocompound metabolism, were also under selection (Supplementary Tables 7g and h).

The allotetraploid nature of bread wheat raised the question of whether the duplicate copies of the homoeologous genes are all under selection. Using the ordered sets of homoeologous genes, we established the syntenic relationships of the selected targets on the other subgenomes (Fig. 4b). This analysis indicated that some syntenic regions shared the signature of selection; for example, the syntenic regions of chromosomes 6A, 6B, and 6D were all under selection (Fig. 4b, Supplementary Table 8i). In the majority of cases however, we could not identify selection signatures in the syntenic regions, suggesting that the homoeologous genes are generally under differing selection pressures and are likely responsible for new functions.

**Fig. 4.**
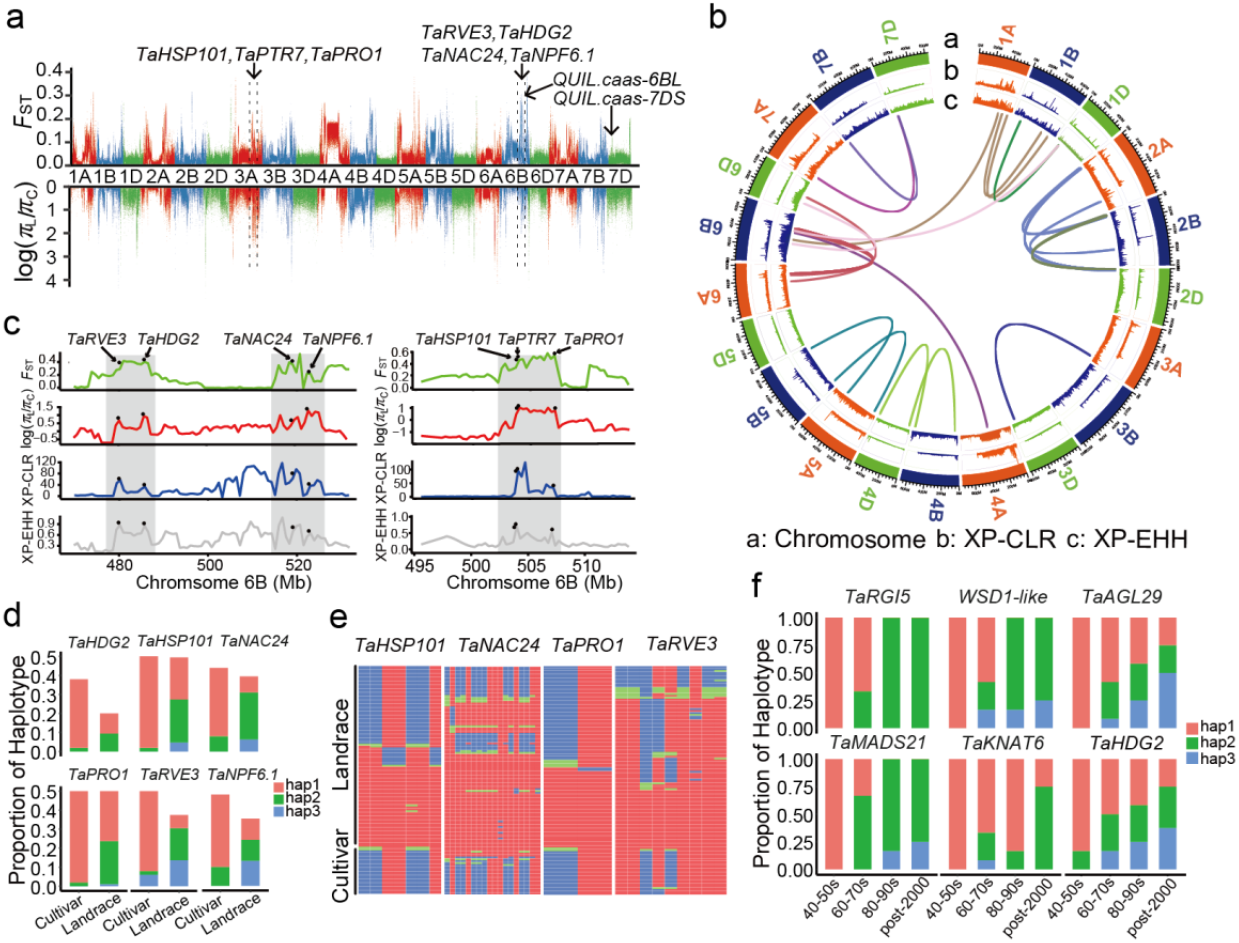
Population sweep selection during modern breeding between the cultivars and landraces. **(a)** The distribution of the *F*_ST_ and θ_π_ ratio (landrace/cultivar) along the chromosomes. The genome-wide threshold was defined by the top 5% of the *F*_ST_ and θ_π_ ratio (landrace/cultivar) values. Candidate genes and QTLs were marked on the map. **(b)** The distribution of XP-CLR and XP-EHH scores along the chromosomes. From the outside to the inside, the data presented are the XP-CLR score, XP-EHH score, and the duplicated copies of the homoeologous genes in the different subgenomes under selection for every chromosome. **(c)** Seven genes distributed in three candidate regions were commonly found to be under selection using the *F*_st_, θ_π_ ratio, XP-CLR, and XP-EHH methods. The black points indicate the location of the candidate selected genes on the chromosomes. **(d)** The haplotype frequency of the candidate genes between the Chinese landrace and cultivar populations. **(e)** Heatmap of genotypes in the putative selective sweeps. **(f)** The selection of the favored haplotype frequencies in modern Chinese cultivars during wheat breeding since the 1940s.

To identify the high-confidence sweep-selected regions, we focused on the overlapping regions commonly identified using the three above-mentioned approaches, which contained 48 HC genes (Supplementary Table 8e). Chromosome 6B had the strongest selection signal. A sub-region of this selected region contains *TaNPF6.1-6B* (Fig. 4c), an ortholog of the *Arabidopsis* nitrate transporter gene NRT1.1^37,38^. Within the same selected region, we also found *TaNAC24*, the ortholog of a gene that confers heat and drought tolerance in rice^39^. In a neighboring subregion, we found strong selection signals for *TaRVE3* (which encodes a putative circadian clock and flowering time regulator)^40^, and *TaHDG2* (encoding a putative stomata and epidermis patterning regulator)^41^, suggesting the existence of a selected region regulating multiple traits on chromosome 6B (Fig. 4c). In another selected region on chromosome 3A, we identified *TaHSP101*, which enhances tolerance to salt and desiccation stresses^42^; *TaPTR7*, the ortholog of a rice NAC transcription factor conferring salt tolerance^43^; and *TaPROl*, a component on the actin cytoskeleton potentially related to grain size^44^ (Fig. 4c). The majority of the selected features and genes are currently poorly annotated and warrant further investigation.

We analyzed the haplotype frequency of the annotated genes under sweep selection, except *TaPTR7* as it contained no SNPs. In the landraces, the frequencies of the primary haplotypes of all selected genes were lower than in the cultivars (Fig. 4d and 4e). Different periods of wheat breeding history likely had diverse breeding goals. To clarify the changes in the haplotypes of these genes over time, we screened the SNPs of six genes in the selected regions *(TaAGL29, TaHDG2, TaKNAT6, TaMADS21, TaRGI5*, and *WSD1-like)* for Chinese cultivars released between the 1940s and the 2000s (Supplementary Table 12b). For each gene, the frequency of the favorable haplotype gradually increased over time (Fig. 4f).

### Local adaption and related traits

Our phylogenetic analysis showed that landraces from geographically close regions tended to be more genetically similar (Fig. 2). To understand the selection undergone by the genes and corresponding traits of these genotypes during geographic diversification and/or local breeding, we calculated the population differentiation across different geographic groups.

The European landraces contributed the majority of the A and B subgenomes of the Chinese cultivars (Supplementary Figs. 7 and 8). We compared these two groups to identify the regions selected during the local adaptation of the Chinese cultivars. Using the *F*_ST_-θ_π_ approach, we identified 13.08 Mb of putatively selected regions covering 729 genes (Fig. 5a). Over a dozen regions were similarly selected in two or three of the subgenomes (Fig. 5b). We also applied the XP-EHH and XP-CLR methods, which led to the identification of a 2.68-Mb putatively selected region (covering 951 genes) and a 26.08-Mb putatively selected region (covering 3,232 genes), respectively (Supplementary Table 9). Finally, we excluded the selected regions identified in our earlier comparison of all cultivars and landraces to focus on local adaption. A total of 3,814 genes were identified using all three methods, with 51 genes being commonly detected (Fig. 5d and Supplementary Table 9e). Among the regions commonly identified using all three approaches, we found that *TaPGIP1*, a polygalacturonase-inhibiting defense protein^45^; *WRKY27*, involved in disease resistance^46^; *TaNAK*, a putative protein kinase involved in defense^47^; and *TaPHB2*, a mitochondrial prohibitin complex protein^48^, were all located within a selected region on chromosome 2A (Fig. 5c). The regions on chromosomes 2B and 2D syntenic to this region were also under selection. Another selected region on chromosome 3A contains *TaPIN1*, required for auxin-dependent root branching^49^; *TaPrx*, related to oxidative stress^50^; and *TaPEPR1*, involved in the innate immunity response^51^ (Fig. 5c). A detailed haplotype frequency analysis indicated that the frequencies of the primary haplotypes in the European landraces were lower than in the Chinese cultivars for all six genes (Fig. 5e and f).

**Fig. 5.**
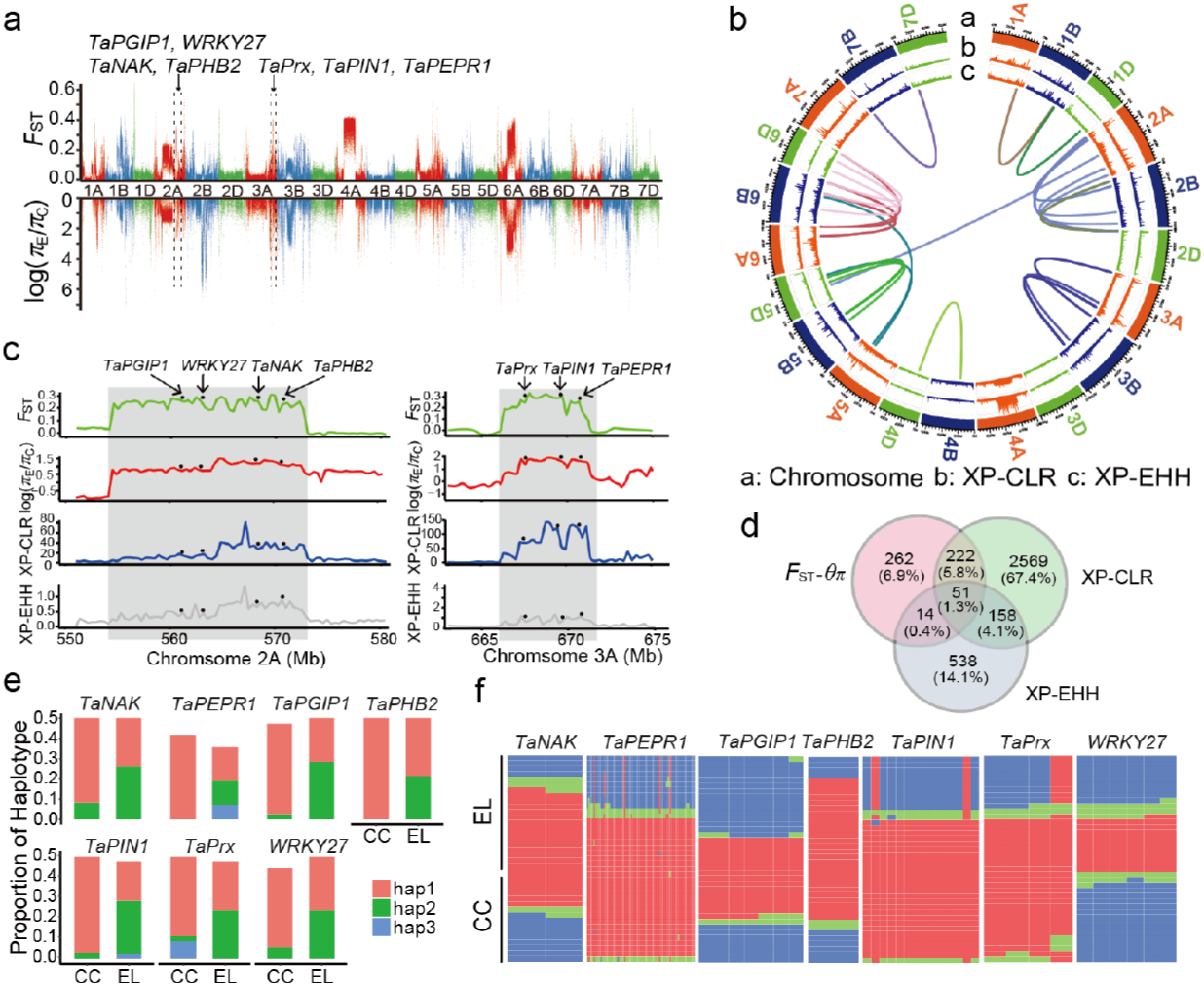
Population sweep selection during local adaption in Chinese cultivars and European landraces. (a) The distribution of the *F*_ST_ and θ_π_ ratio (European landrace/Chinese cultivar) along the chromosomes. The genome-wide threshold was defined by the top 5% of the *F*_ST_ and θ_π_ ratio (European landrace/Chinese cultivar) values. Candidate genes and QTLs are marked on the map. (b) The distribution of XP-CLR and XP-EHH scores along the chromosomes. From the outside to the inside, the data presented are the XP-CLR score, the XP-EHH score, and the duplicated copies of the homoeologous genes in the different subgenomes under selection along every chromosome. (c) Seven genes distributed in the two candidate selection regions identified using the *F*_ST_-θ_π_ ratio, XP-CLR, and XP-EHH methods. The black points indicate the location of the candidate genes on the chromosome; every gene has a strong selection signal. (d) A Venn diagram of wheat genes under selection detected using the *F*_ST_-θ_π_ ratio, XP-CLR, and XP-EHH methods. (e) The haplotype frequency of the candidate genes between the Chinese cultivars (CC) and the European landraces (EL). (f) Heatmap of genotypes in the putative selective sweep.

The West Asian landraces are the most closely related modern relatives of the ancestral wheat population. We therefore compared the European and Chinese landraces to the West Asian landraces to identify the regions selected during local adaptation as wheat gradually spread to the west and the east. Using *F*_ST_-θ_π_, XP-EHH, and XP-CLR analyses, we identified 22.75-Mb (covering 1,760 genes), 3.00-Mb (covering 1,074 genes), and 21.04-Mb (covering 2,619 genes) putatively selected regions between the European and West Asian landraces, respectively (Supplementary Table 10). These selected regions overlap with a few reported QTLs, including two associated with flour quality^52^. We identified a significantly selected region on chromosome 2B that contains *TaNPF6.1-2B*, another ortholog of the *Arabidopsis* nitrate transporter gene *NRT1.1* (Supplementary Fig. 12a). The same region also harbors *TaFBP7*, a gene related to temperature response^53^, and *TaANP1*, an oxidative stress responsive gene^54^. Whereas novel haplotypes for *TaNPF6.1-2B* emerged in the European landraces, the genotypes typically maintained one of the two *TaFBP7* and *TaANP1* haplotypes from the West Asian landraces (Supplementary Fig. 12b and c).

When comparing the Chinese landraces with the West Asian landraces, we identified 8.94 Mb of putatively selected regions (covering 550 genes) using *F*_ST_-θ_π_, 2.81 Mb (covering 1,027 genes) using XP-EHH, and 18.85 Mb (covering 2,392 genes) using XP-CLR (Supplementary Table 11). One of the genes identified in the selected regions was *TaOSH43* (Supplementary Fig. 13a), which regulates shoot meristem homeostasis and thus plant architecture and spike development^55^. A new haplotype emerged and became dominant in the Chinese landraces (Supplementary Fig. 13b and c). We also found that a flour-quality QTL^52^ overlaps with a selected region on chromosome 6B.

## Discussion

Our analysis of 120 representative bread wheat accessions revealed a unique pattern of genome changes that occurred during wheat adaption and modern breeding, especially in China. The allopolyploidization of hexaploid bread wheat is expected to have occurred under human cultivation, as bread wheat only exists in cultivated forms^3,10^. Consistent with this, bread wheat has a substantially lower genome diversity than other crop and weed species (Fig. 1c and Table 1). Nevertheless, gene flow from the allotetraploid *T. turgidum* (AABB) is likely to have contributed to the genetic diversity of the A and B subgenomes in bread wheat^27,28^, which means these two subgenomes harbor more selective sweeps, as was demonstrated here. By contrast, the barrier to gene flow from the diploid *Ae. tauschii* (DD) to bread wheat is much higher, as suggested by the substantially lower nucleotide diversities in the D subgenome, higher numbers of rare mutations, and fewer selective sweeps reported here.

Bread wheat has a long cultivation history and has been adapted to a wide range of environmental conditions. The rich genetic diversity in cultivated accessions and the lack of a wild bread wheat mean the gene flow among geographic regions is potentially substantial. Indeed, we found that modern wheat breeding in China significantly used the genetic diversity of the European landraces. In future breeding programs, the genetic diversity of landraces from diverse geographic regions may provide the additional genetic diversity required to fulfill the demands of our increasing population and changing climate.

We found that different ancestors made distinct contributions to each subgenome. In the Chinese cultivars, the genetic diversity of the A and B subgenomes was mainly contributed by the European landraces. By contrast, the D subgenome of the Chinese cultivars contains more diversity from local landraces. Although the European landraces and the Chinese landraces have similar levels of genetic diversity for each subgenome (Fig. 2d), the European landraces experienced continuous gene exchange with the allotetraploid *T. turgidum.* Such a gene exchanges may diversify favorable alleles that would be selected during modem breeding in China. On the other hand, the D subgenome has limited genetic exchange in all landraces. As a result, the D subgenome of the European landraces may not have additional advantages over Chinese landraces as the A and B subgenomes have. These differences may explain the different contribution of AB and D subgenomes to the Chinese cultivars. Alternatively, the selection of accessions may lead to inevitable bias, although we have used genome-wide markers to ensure that the resequenced accessions represent the widest genome diversity.

We have narrowed down the selective sweeps corresponding to selection during wheat diversification and modern breeding, which will be useful for the future characterization of genes underlying important traits. Consistent with the contributions of different ancestors to each subgenome, we found that the homoeologous genes in the different subgenomes are often under differential selection, suggesting that it may be feasible to target a single copy of a homoeologous gene during breeding.

## Methods

### Plant materials

A total of 120 hexaploid wheat *(Triticum aestivum)* accessions from Asia, Europe, and America were used in this study, including 95 landraces and 25 cultivars. Among them were 60 accessions, including 42 landraces, and 18 elite varieties, from a Chinese mini core collection, which is estimated to represent the majority of the genomic diversity (~70%) of the 23,090 accessions in the Chinese national collection^16,56^. Based on genome-wide simple sequence repeat (SSR) data, these 60 accessions were estimated to represent the widest genome diversity of the mini core collection. The rest of the accessions were selected following the sequencing of 326 accessions collected from major wheat cultivation sites using the genotyping-by-sequencing (GBS) method^57^. Based on the GBS results, the other 60 varieties were selected for inclusion in the study, including 16 from West Asia, 12 from Central and South Asia, 1 from Japan, 24 from Europe, and 7 from the Americas, which represent the widest genome diversity. All the accessions were purified in multiple rounds of single-seed descent to ensure their homozygosity.

### Library preparation for Illumina next-generation sequencing

Approximately 1.5 μg genomic DNA was extracted from each sample using the CTAB method and prepared in libraries for sequencing using a TruSeq Nano DNA HT sample preparation kit (Illumina, USA) following the manufacturer’s recommendations. Briefly, the genomic DNA samples were fragmented by sonication to a size of ~350 bp, then end-polished, A-tailed, and ligated with the full-length adapters for Illumina sequencing with further PCR amplification. Index codes were added facilitate the differentiation of sequences from each sample. The PCR products were purified (AMPure XP bead system; Beckman Coulter, USA) and the libraries were analyzed for their size distribution using an Agilent 2100 Bioanalyzer (Agilent Technologies, USA) and quantified using real-time PCR.

### Genome sequencing and quality control

The libraries were sequenced using the Illumina HiSeq X platform (Illumina). In total, ~21,676 Tb of raw data were generated. To ensure reliable reads without artificial bias, the low-quality paired reads (≥ 10% unidentified nucleotides (N); > 10 nucleotides aligned to the adaptor, allowing ≤ 10% mismatches; > 50% bases with a Phred quality less than 5) were removed. A total of 21,531 Tb (~179.4 Gb per sample) high-quality genomic data was obtained.

### Read mapping and SNP calling

The remaining high-quality paired-end reads were mapped to the bread wheat reference genome (IWGSC RefSeq v1.0) using Burrows-Wheeler Aligner software with the command ‘mem -t 4 -k 32 –M’^58^. To reduce mismatches generated by the PCR amplification before sequencing, the duplicated reads were removed with the help of SAMtools (v0.1.19)^59^. After the alignment, SNP calling was performed on a population scale using a Bayesian approach in SAMtools. The genotype likelihoods were calculated from the reads of each individual at each genomic location, and the allele frequencies were determined using a Bayesian approach. The ‘mpileup’ command was used to identify SNPs using the parameters ‘-q 1 -C 50 -S -D -m 2 -F 0.002’. To exclude the SNP calling errors caused by incorrect mapping or InDels, only the 70,172,660 high-quality filtered SNPs (depth ≥ 4, maf ≥ 0.01, miss ≤ 0.1) were used in the subsequent analysis.

### Functional annotation of genetic variants

The SNP annotation was performed using the published wheat genome^9^ and the ANNOVAR package (v2013-05-20)^60^. Based on the genome annotation, the SNPs were categorized as being in exonic regions (overlapping with a coding exon), intronic regions (overlapping with an intron), splicing sites (within 2 bp of a splicing junction), upstream or downstream regions (within a 1-kb region upstream or downstream of a transcription start or stop site, respectively), or intergenic regions. The SNPs in the coding exons were further grouped into synonymous SNPs (do not cause amino acid changes) or non-synonymous SNPs (cause amino acid changes). The mutations causing stop gain and stop loss were also classified into this group. To exclude genes with possible structural annotation errors, those expressed in the leaves or existing in multiple copies were selected as HC genes for further analysis.

### Population genetic diversity

Nucleotide diversity θπ and fixation index (*F*_ST_) were calculated using VCFtools (v0.1.14)^61^, while Watterson’s estimator (θ_w_) and Tajima’s *D* were calculated using VariScan (v2.0.3)^62^. These population statistics were analyzed using the sliding-window approach (20-kb windows with 10-kb increments). HC genes were selected to identify single-copy orthologous genes between the A, B, and D subgenomes using Proteinortho (v5.16), with default settings^63^.

### Linkage disequilibrium analysis

To estimate and compare the patterns of linkage disequilibrium (LD) between different populations, the squared correlation coefficient (r^2^) between pairwise SNPs was computed using the software Haploview (v4.2)^64^. The program parameters were set as ‘-n -dprime -minMAF 0.1’. The average *r*^2^ value was calculated for pairwise markers in a 500-kb window and averaged across the whole genome.

### Phylogenetic tree and population structure

A total of 1,925,854 SNPs in coding regions (exonic and intronic) were used for the population genetics analysis. To clarify the phylogenetic relationship from a genome-wide perspective, an individual-based neighbor-joining tree was constructed using the *p*-distance in the software TreeBeST (v1.9.2), with bootstrap values determined from 1000 replicates^65^.

The population genetic structure was examined using the program ADMIXTURE (v1.23)^66^. First, 95 landraces were used to estimate the genetic ancestry, specifying a *K* ranging from 2 to 8. The most suitable number of ancestral populations was determined to be *K* = 3, for which the lowest cross-validation error of 0. 492 was obtained. A principal component analysis (PCA) was also conducted to evaluate the genetic structure of the populations using the software GCTA^67^. First, a genetic relationship matrix (GRM) was obtained using the parameter ‘–make-grm’, then the top three principal components were estimated with the parameter ‘–pca3’.

### Population admixture analyses

The population relatedness and migration events were inferred using TreeMix^68^. A total of 113 varieties with little admixture were selected to represent the seven subgroups. The 1,925,854 coding-region SNPs were used to build a maximum likelihood tree, using a window size of 2000 SNPs. This was repeated 10 times, using the West Asian landraces as the root group. The tree with the lowest standard error for the residuals was selected as the base tree topology. The population pairs with an above zero standard residual error were identified as candidates for admixture events, which represent populations which the data indicate are more closely related to each other than is demonstrated in the best-fit tree^68^. TreeMix was then run using between one and six introduced migration events. When three migration events were added, the residuals were much lower than for the trees generated using other numbers of migration events.

### Introgression analysis

The identity scores (IS) were calculated^32^ to visualize the shared haplotypes between the three populations (European varieties, Chinese landraces, and Chinese cultivars). The IS values were used to evaluate the similarities of every sequenced sample to the reference genome (Chinese Spring) within 20-kb windows. The IS was calculated using the following formula:

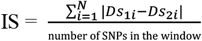

where *D* represents genotype similarity to reference genome of samples in single site. The IS of any single site was calculated as the difference in the *D* between two samples. The average IS value was calculated for each population.

The *H*_P_ (population heterozygosity) was calculated for each candidate introgression region. At each detected SNP position, the numbers of reads corresponding to the most and least frequently observed allele (nMAJ and nMIN, respectively) were counted in each population. The H_P_ for each window was calculated using the following formula:

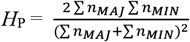

Σ*n*_MAJ_ and Σ*n_MIN_* are sums of the *n_MAJ_* and *n_MIN_* values, respectively, calculated for all SNPs in the 20-kb window (sliding in 10-kb steps)^69^. The population recombination rate was estimated from the SNPs of three populations (European varieties, Chinese landraces, and Chinese cultivars) within a 20-kb window using the R package FastEPRR (v1.0)^70^.

### Genome-wide selective sweep analysis

A sliding-window approach (20-kb windows sliding in 10-kb steps) was applied to quantify the levels of polymorphism (*θ_π_*, the pairwise nucleotide variation as a measure of variability) and genetic differentiation (*F*_ST_) between the different populations using the VCFtools software (v0.1.14)^61^. The *θπ* ratios were log_2_-transformed. Subsequently, the empirical percentiles of the *F*_ST_ and log_2_(*θ_π_* ratio) values in each window were estimated and ranked. The windows with the top 5% *F*_ST_ and log_2_(*θ_π_* ratio) values were considered simultaneously as candidate outliers under strong selective sweeps. All outlier windows were assigned to corresponding regions and genes. The cross-population extended haplotype homozygosity (XP-EHH) statistic was estimated^71^ for the cultivar group and the landrace group, using the landrace group as a contrast. The genetic map was assumed to be 0.18 cM/Mb for the A and B subgenome, and 0.341 cM/Mb for the D subgenome. The XP-CLR score^35^ was used to confirm the selective sweeps on the basis of domestication features, with the highest being 1%, via the cross-population composite likelihood method.

### Candidate gene analysis

Gene Ontology (GO) and Kyoto Encyclopedia of Genes and Genomes (KEGG) analyses were performed on the candidate genes (all annotated genes in the outlier windows denoted by the top 5% of the *F*_ST_ and log2θ ratio) values, the top 1% of the XP-EHH values, and the top 1% of the XP-CLR scores). The candidate genes were also attributed to known KEGG pathways (http://www.kegg.jp).

### PCR primers and amplicon sequence analysis

A total of 28 Chinese cultivars developed between the 1940s and the 2000s were analyzed to clarify the changes in the haplotypes of the candidate genes during wheat breeding. All primers were designed using the Primer 5.0 software and listed in Table S13a. DNAMAN8.0 was used to align the sequences to the reference genome and identify the SNPs, after which the frequencies of the main haplotypes in the candidate genes were analyzed.

## Data availability

The sequencing data from this study have been deposited into the Sequence Read Archive (https://www.ncbi.nlm.nih.gov/sra) under accession PRJNA439156.

## Acknowledgments

We thank Yalong Guo and Wenfeng Qian for their valuable suggestions. This research was supported by the Chinese Academy of Sciences (CAS; grant no. XDA08020105 to Y.J. and XDA08040108 to H.C.), the National Natural Science Foundation of China (grant no. 31430010 to Y.J., 31871245 to Ying W, and 31770311 to C.T.), the National Program for Support of Top-Notch Young Professionals (Y.J.), University of CAS (grant no. 110601M206 to Ying W.), the Beijing NOVA Program (grant no. Z161100004916107 to Ying W.), the National Transgenic Science and Technology Program (grant no. 2016ZX08010-002 to C.T.), and the CAS Youth Innovation Promotion Association (grant no. 2017139 to C.T.).

## Author information

These authors contributed equally: H. Chen, C. Jiao, Ying Wang, Yuange Wang.

## Affiliantions

*State Key Laboratory of Plant Genomics, Institute of Genetics and Developmental*

*Biology, Chinese Academy of Sciences, Beijing 100101, China*

Haofeng Chen, Yuange Wang, Caihuan Tian, Haopeng Yu, Jing Wang, Yuling Jiao

*Novogene Bioinformatics Institute, Beijing 100083, China*

Chengzhi Jiao, Wenkai Jiang, Hongfeng Lu

*College of Life Sciences, University of Chinese Academy of Sciences, Beijing 100049, China*

Ying Wang, Fei Lu, Xiangdong Fu, Yongbiao Xue, Hongqing Ling, Yuling Jiao

*West China Biomedical Big Data Center, West China Hospital/West China School of Medicine, and Medical Big Data Center, Sichuan University, Chengdu 610041, China*

Haopeng Yu

*Department of Crop Genomics and Bioinformatics, College of Agronomy and Biotechnology, National Maize Improvement Center of China, China Agricultural University, Beijing 100193, China*

Xiangfeng Wang

*State Key Laboratory of Plant Cell and Chromosome Engineering, Institute of Genetics and Developmental Biology, Chinese Academy of Sciences, Beijing 100101, China*

Fei Lu, Xiangdong Fu, Yongbiao Xue, Hongqing Ling

## Contributions

Y.J., H.Lu and Ying W. conceived of and designed the study. C.J., Yuange W., H.Y., J.W., X.W., W.J., and H.Lu performed the analyses. C.J., H.Lu, Y.J., Ying W., W.J., and H.Ling interpreted the data. H.C. Yuange W., C.T., J.W. F.L., X.F., Y.X., H.Ling, and Y.J. contributed to data collection. Y.J., C.J., and H.Lu wrote the manuscript with input from all coauthors.

