## Supplemental Figures for "Comparative Population Genomics of Bread Wheat *(Triticum aestivum)* Reveals Its Cultivation and Breeding History in China"

**This PDF file includes:**

Supplementary Figures 1 to 13

Captions for Supplementary Tables 1 to 12

**Other Supplementary information for this manuscript include the following:**

Supplementary Tables 1 to 12 in an Excel file

**Supplementary Figures 1 – 13**

**Supplementary Fig. 1.** Diversity analysis of sequenced wheat accessions.

**Supplementary Fig. 2.** Gene structure annotation of the SNPs in the A, B, and D subgenomes.

**Supplementary Fig. 3.** Linkage disequilibrium (LD; $r^2$) in different chromosomes.

**Supplementary Fig. 4.** The origin and migration route of bread wheat according to Feldmann, 2001.

**Supplementary Fig. 5.** Distribution and population structure ($K$ = 5) of the 95 wheat landraces.

**Supplementary Fig. 6.** TreeMix analysis of the entire wheat genome.

**Supplementary Fig. 7.** Population structure of the A subgenome.

**Supplementary Fig. 8.** Population structure of the B subgenome.

**Supplementary Fig. 9.** Population structure of the D subgenome.

**Supplementary Fig. 10.** Heatmap of the genomic similarity of three wheat populations (European varieties, Chinese cultivars, and Chinese landraces).

**Supplementary Fig. 11.** The statistics of similar genomic regions revealed using the IS method.

**Supplementary Fig. 12.** Population sweep selection between the European and West Asian landraces during their local adaption.

**Supplementary Fig. 13.** Population sweep selection between the Chinese landraces and West Asian landraces during their local adaption.

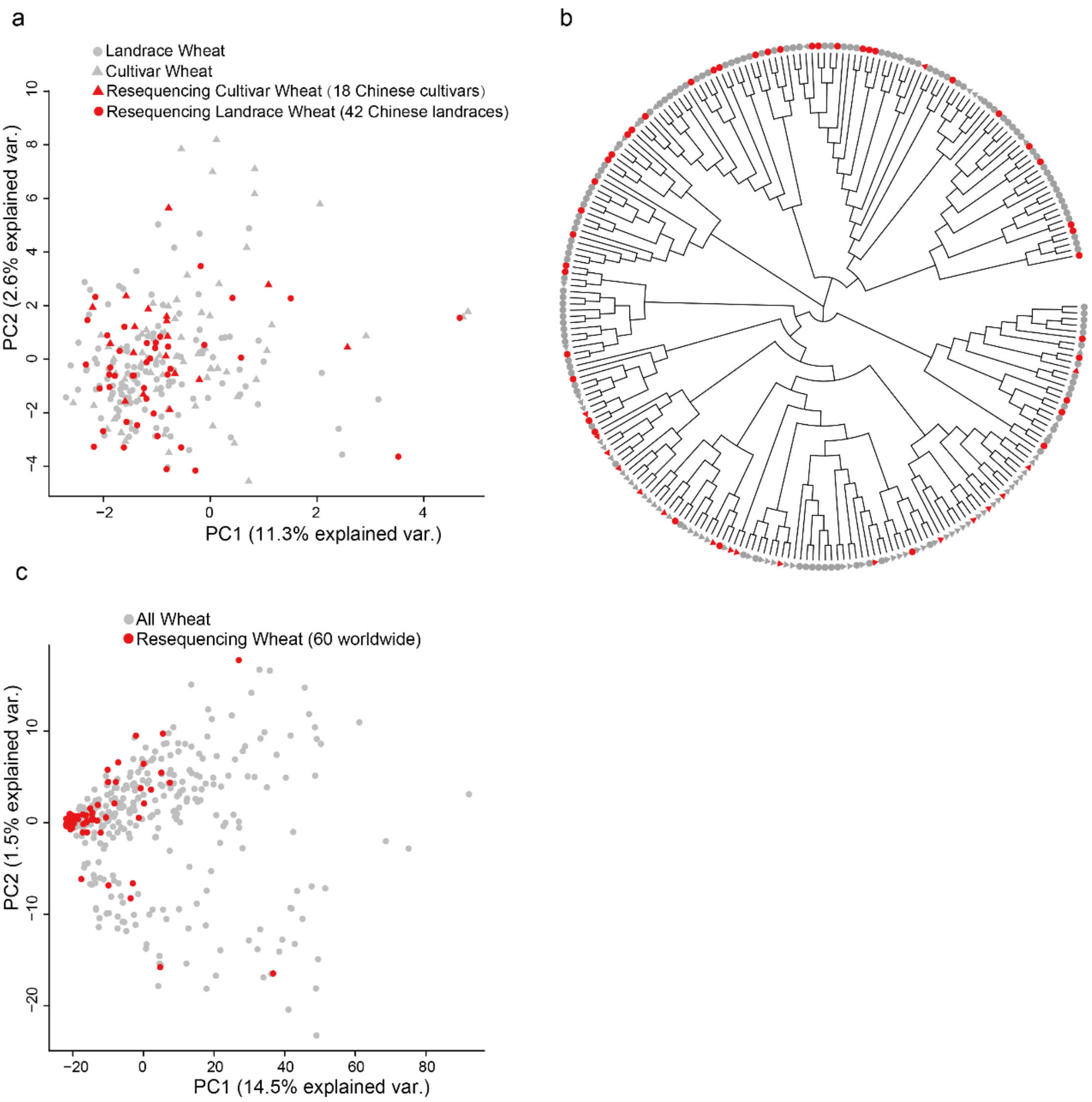


**Supplementary Figure 1**

**Diversity analysis of sequenced wheat accessions.**

(**a**) Principal component analysis of accessions from a Chinese mini core collection 1,2. Accessions selected for resequencing are shown in red; and cultivar and landrace wheat accessions are shown by triangles and circles, respectively. (**b**) Phylogenetic tree of the Chinese mini core collection accessions. Same color code is used as for (**a**). (**c**) Principle component analysis of 326 representative accessions from worldwide with GBS data. Accessions selected for resequencing are shown in red.

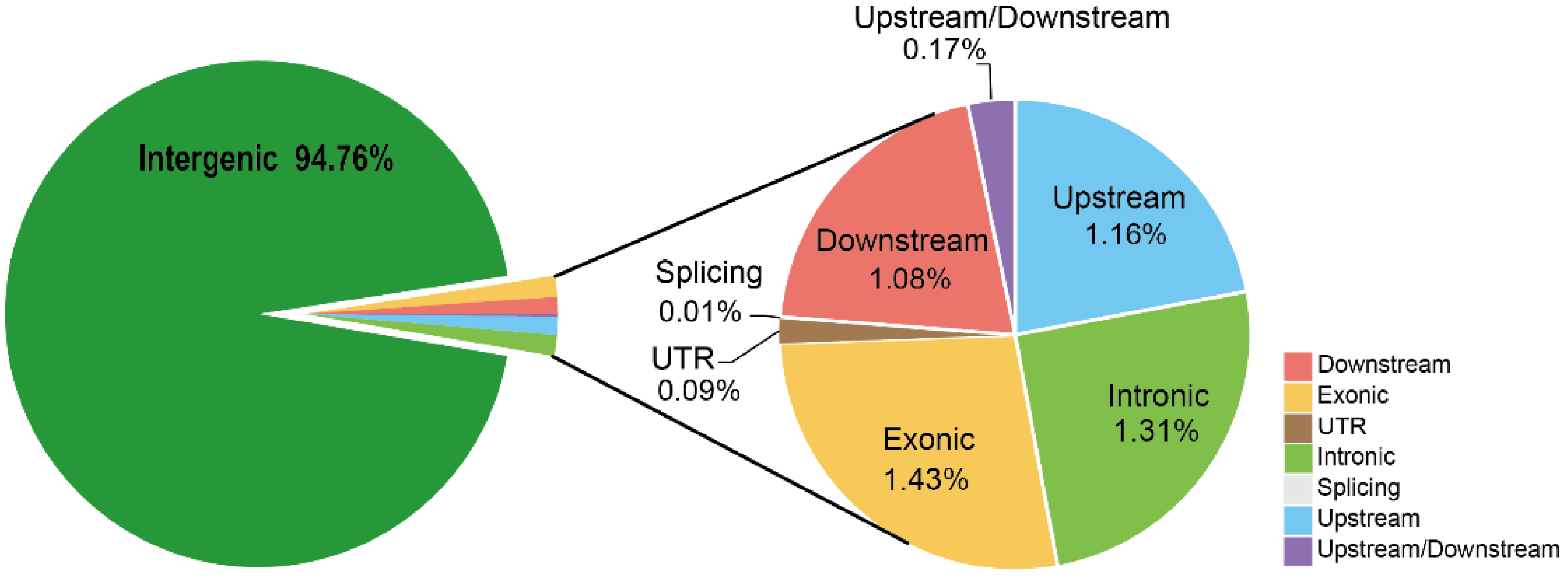


**Supplementary Figure 2**

**Gene structure annotation of the SNPs in the A, B, and D subgenomes.**

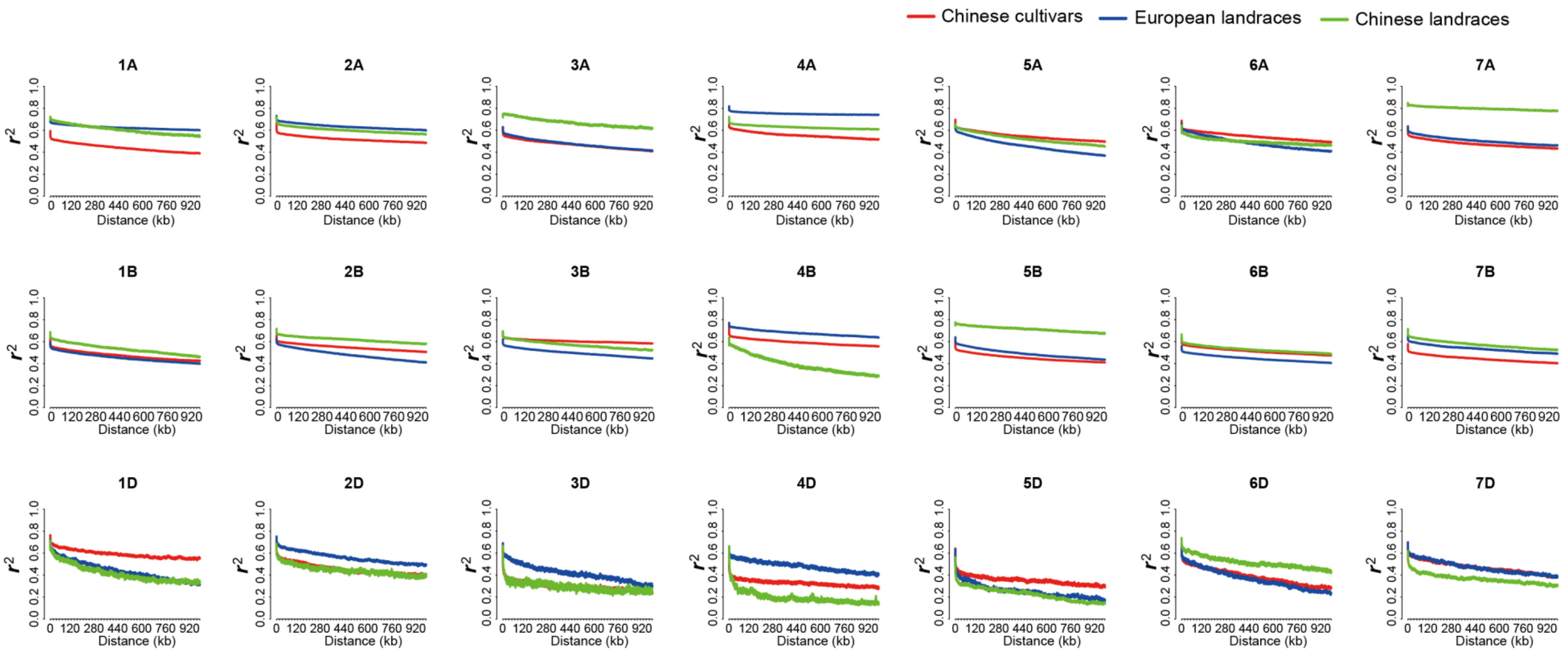


**Supplementary Figure 3**

**Linkage disequilibrium (LD; $r^2$) in different chromosomes.**

The LD decay patterns are presented for three populations (Chinese cultivars, Chinese landraces, and European landraces).

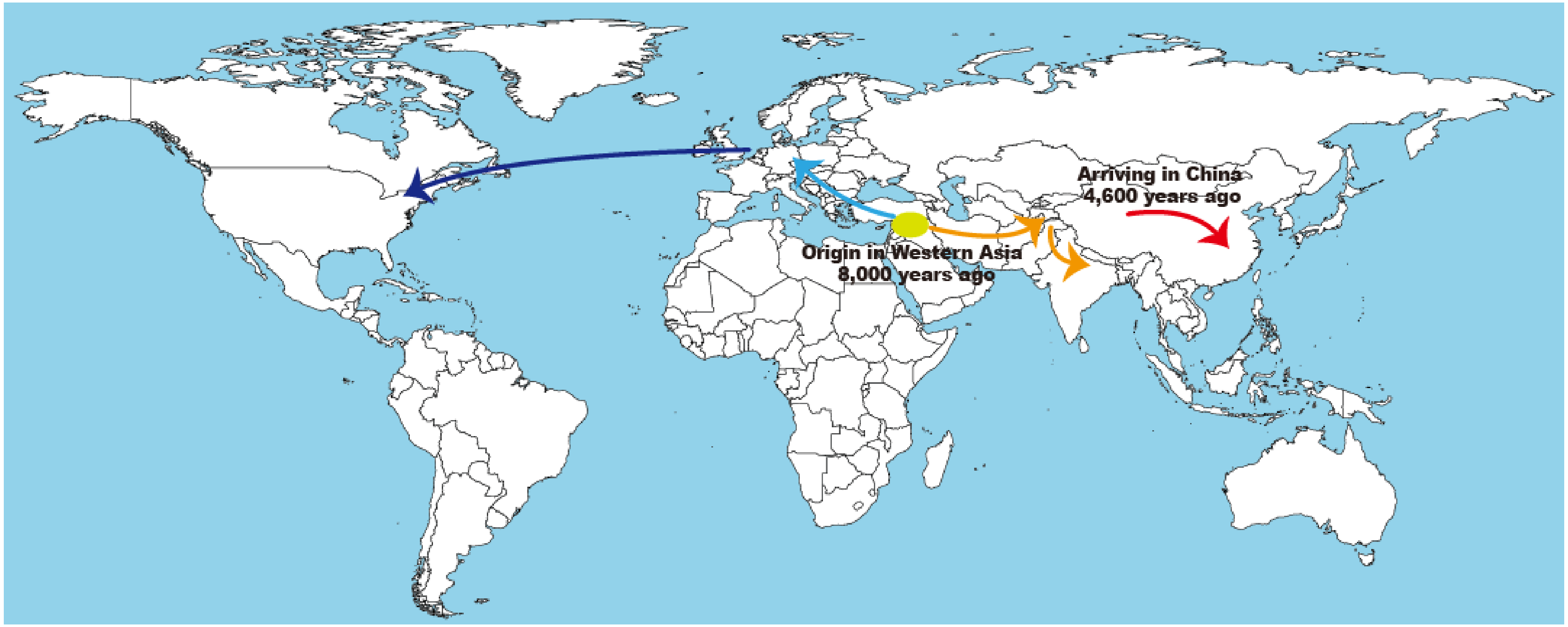


**Supplementary Figure 4**

**The origin and migration route of bread wheat according to Feldmann, 2001.**

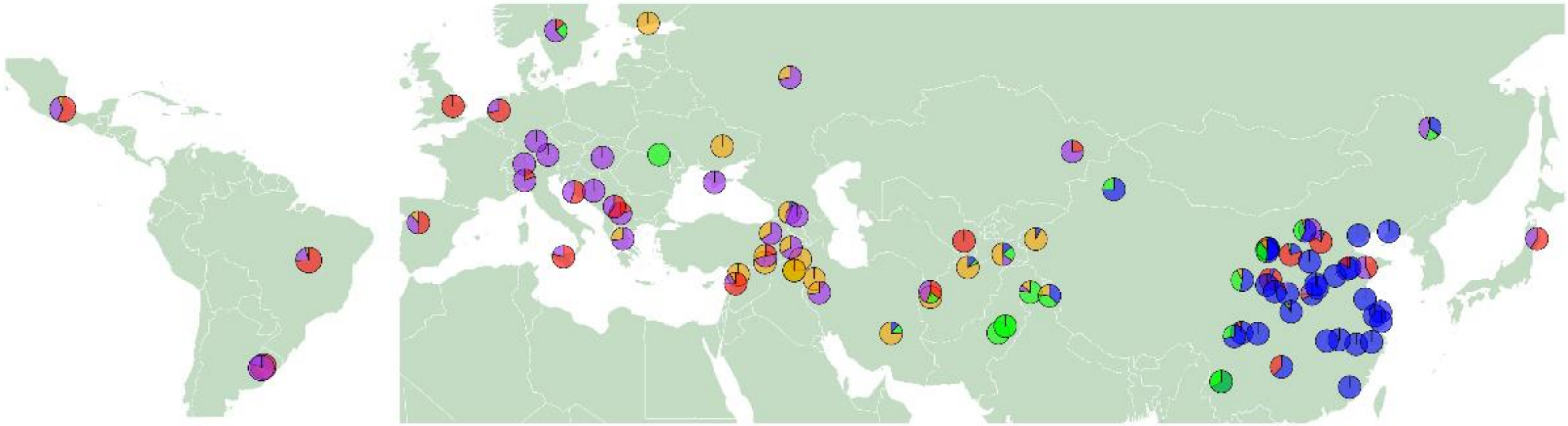

**Supplementary Figure 5**

**Distribution and population structure (*K* = 5) of the 95 wheat landraces.**

The pie charts indicate the proportion of genetic ancestry for *K* = 5, inferred using Admixture. The same color code as in Fig. 2c is used.

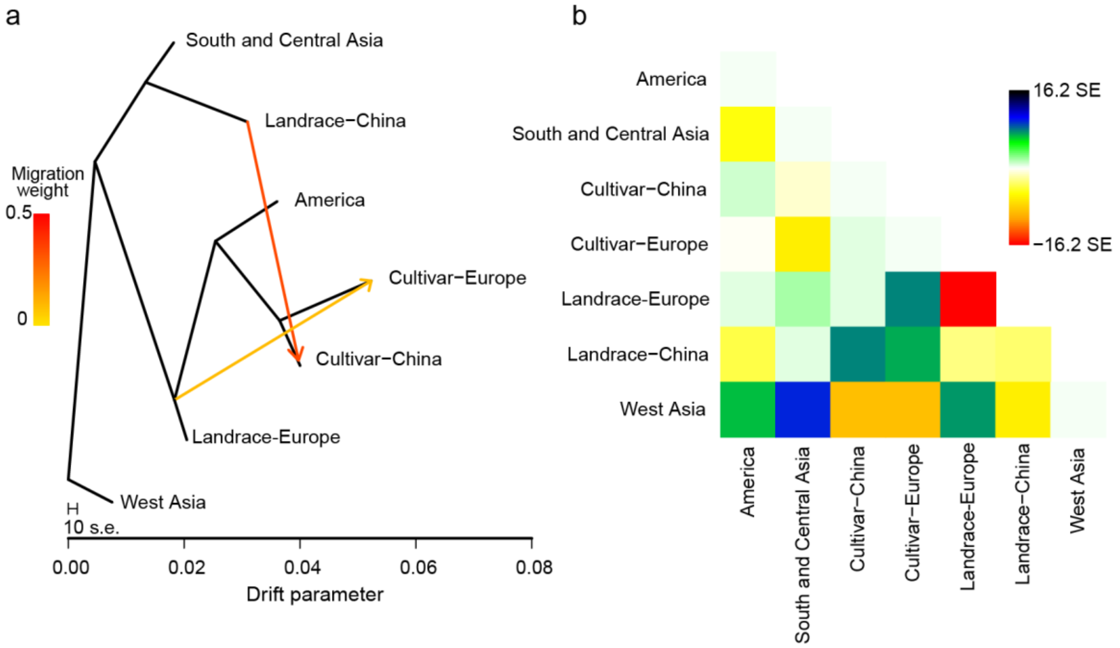


**Supplementary Figure 6**

**TreeMix analysis of the entire wheat genome.**

(**a**) Gene flow between seven populations of wheat. (**b**) Heat map of the gene flow between populations.

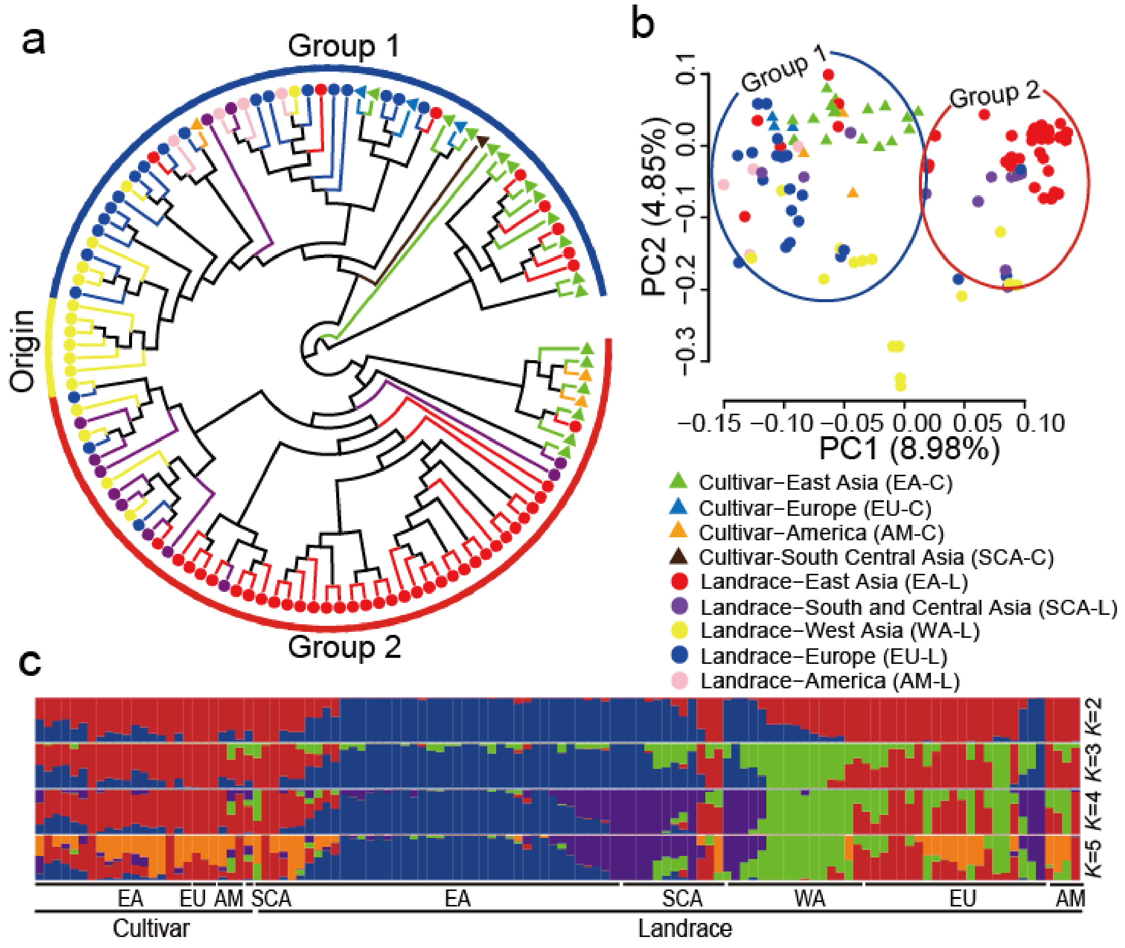


**Supplementary Figure 7**

**Population structure of the A subgenome.**

(**a**) Phylogenetic tree of the A subgenomes in each accession. Triangles represent cultivars, circles represent landraces. Different colors represent populations with different geographical distributions. (**b**) Principal component analysis of the A subgenomes of the wheat accessions. A and B share the same key. (**c**) Population structure of the A subgenomes, determined using Admixture.

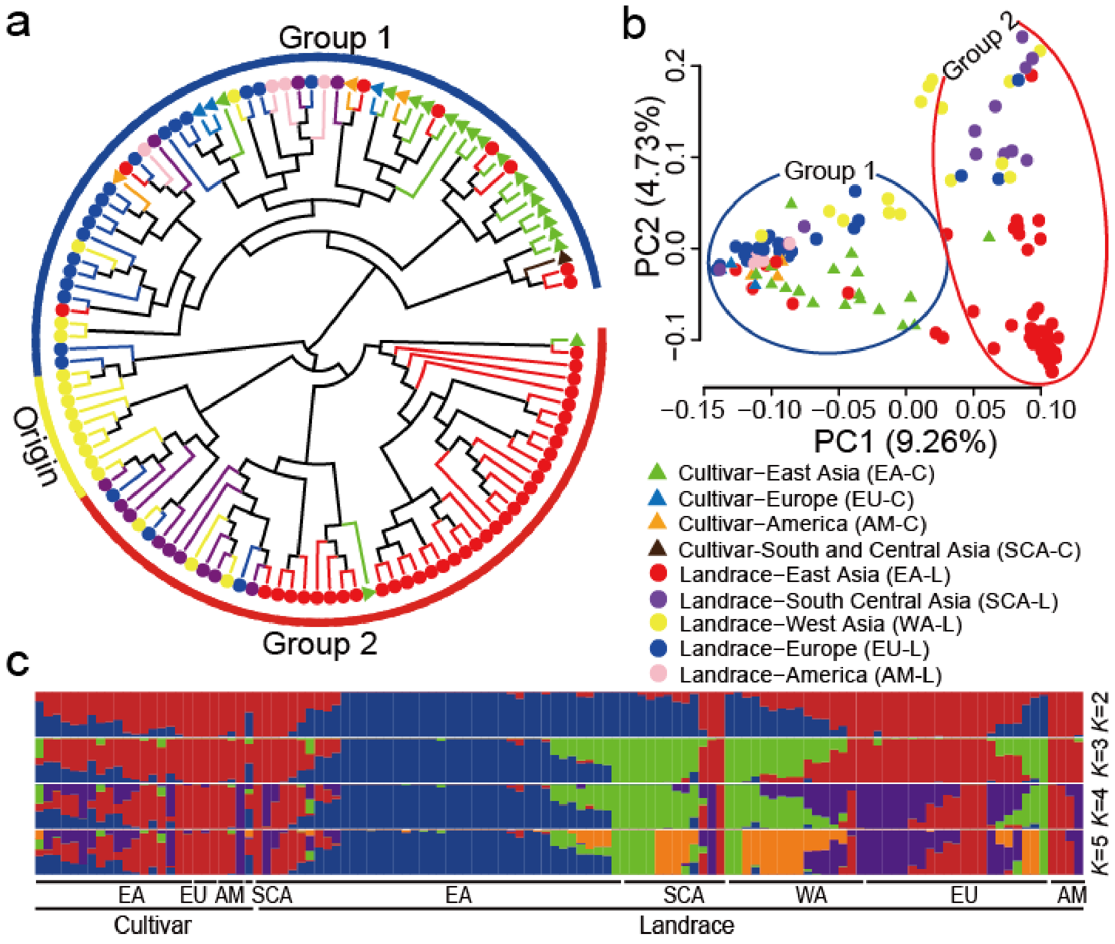


**Supplementary Figure 8**

**Population structure of the B subgenome.**

(**a**) Phylogenetic tree of the B subgenomes in each accession. Triangles represent cultivars, circles represent landraces. Different colors represent populations with different geographical distributions. (**b**) Principal component analysis of the B subgenomes of the wheat accessions. A and B share the same key. (**c**) Population structure of the B subgenomes, determined using Admixture.

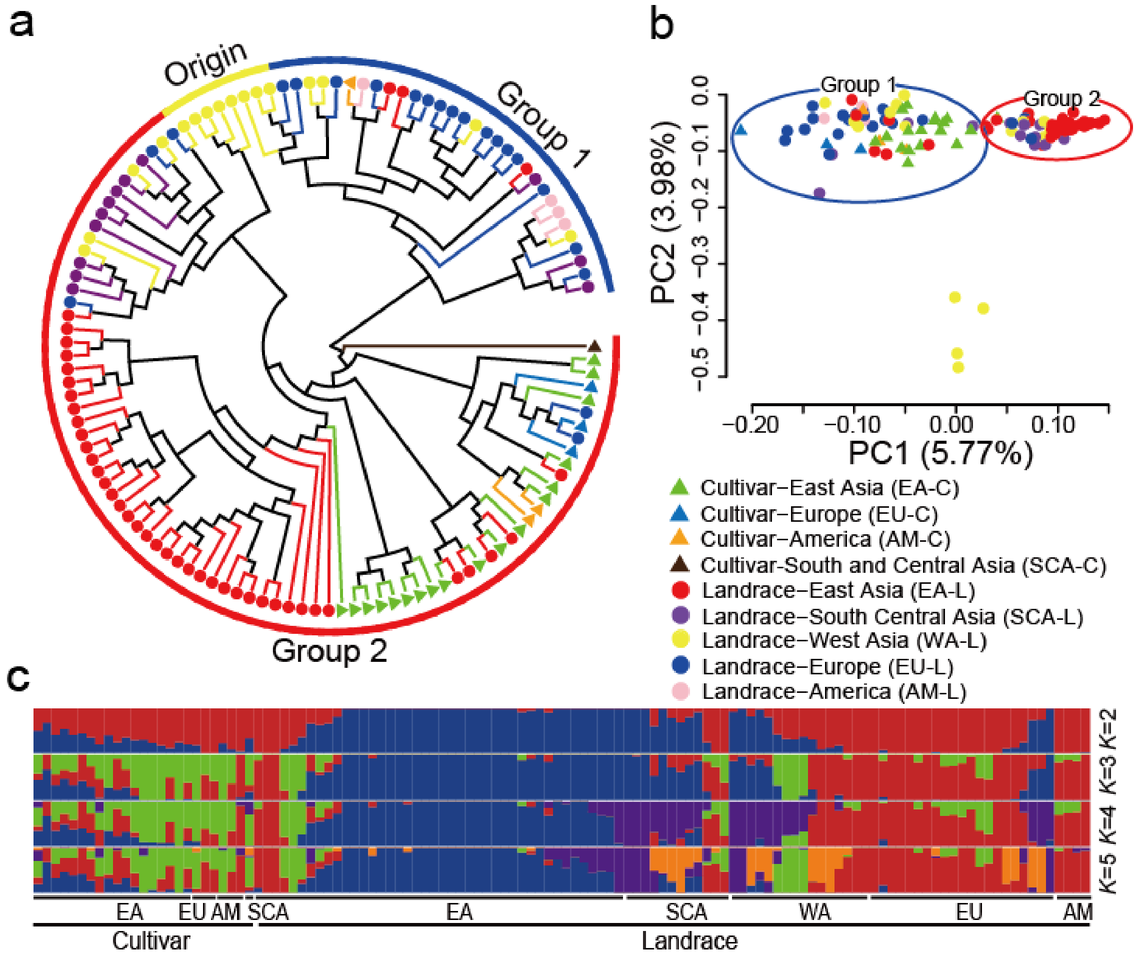


**Supplementary Figure 9**

**Population structure of the D subgenome.**

(**a**) Phylogenetic tree of the D subgenomes in each accession. Triangles represent cultivars, circles represent landraces. Different colors represent populations with different geographical distributions. (**b**) Principal component analysis of the D subgenomes of the wheat accessions. A and B share the same key. (**c**) Population structures of the D subgenomes determined using Admixture.

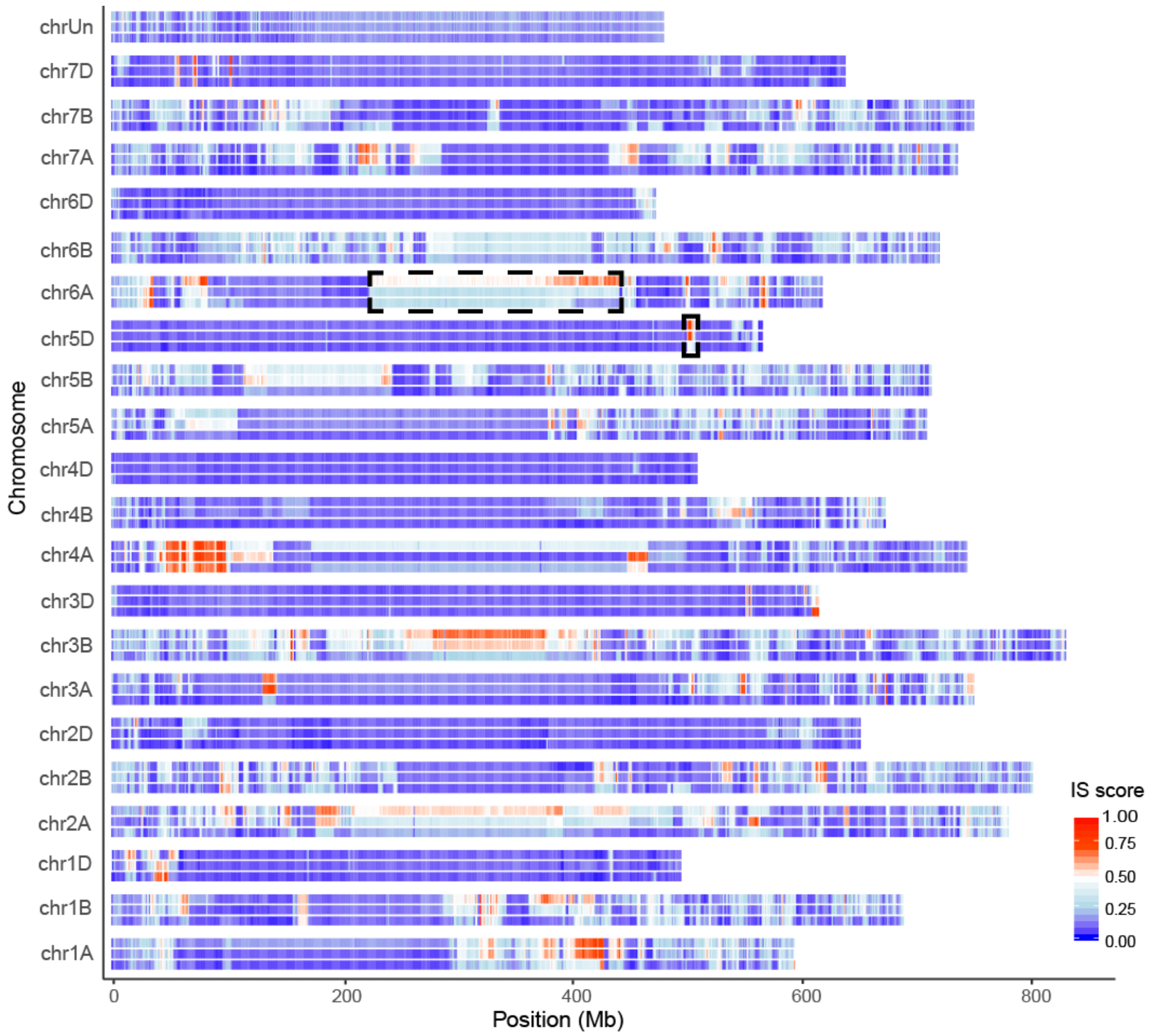


**Supplementary Figure 10**

**Heatmap of the genomic similarity of three wheat populations (European varieties, Chinese cultivars, and Chinese landraces).**

The similarity to the reference genome is shown for each of the 21 chromosomes, in addition to the scaffolds named chrUn. For each chromosome, the heatmap of the IS distribution of the three populations is shown for (from top to bottom) the European varieties, Chinese cultivars, and Chinese landraces. The regions marked with black dotted lines represent putative introgression regions; the one on chromosome 5D is a putative introgression from the European varieties to the Chinese cultivars, while the one on chromosome 6A is a putative introgression from the Chinese landraces to the Chinese cultivars.

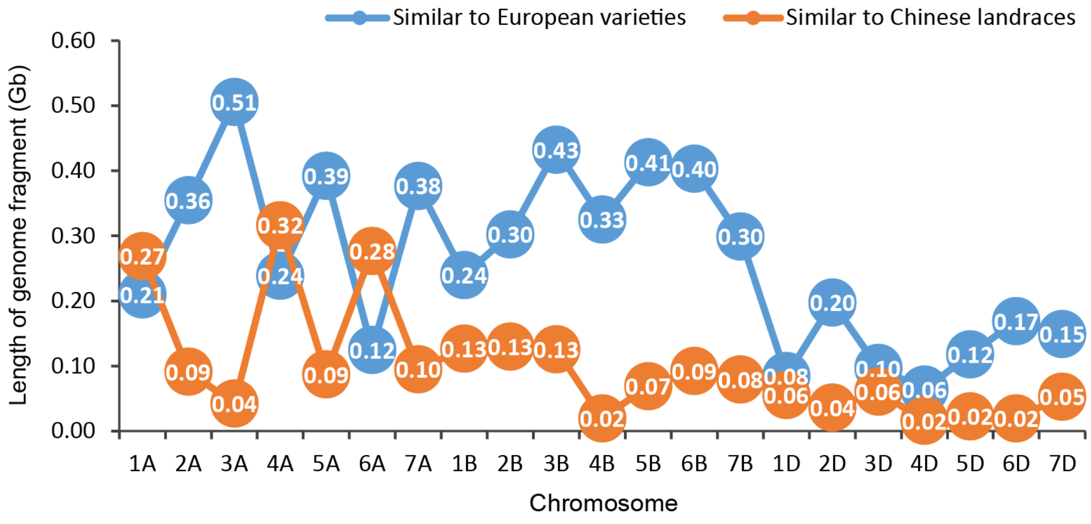


**Supplementary Figure 11**

**The statistics of similar genomic regions revealed using the IS method.**

Each of the chromosomes of the European varieties and the Chinese landraces were compared to those of the Chinese cultivars using the IS methods.

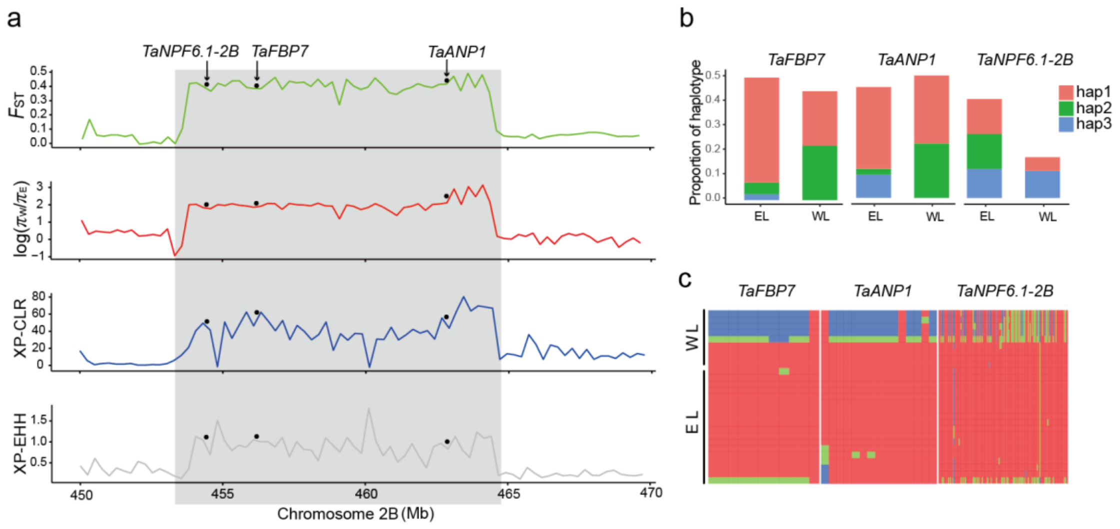


**Supplementary Figure 12**

**Population sweep selection between the European and West Asian landraces during their local adaption.**

(**a**) Three genes distributed in a candidate region were commonly identified using the $F_{ST}$-$\theta\pi$ ratio, XP-CLR, and XP-EHH methods. The black points indicate the location of these genes, each of which had a strong selection signal. (**b**) The haplotype frequencies of the candidate genes between the European landraces (EL) and West Asian landraces (WL). (**c**) Heatmap of the European and West Asian landrace genotypes in the putative selective sweeps containing *TaNPF6.1-2B*, *TaFBP7*, and *TaANP1*.

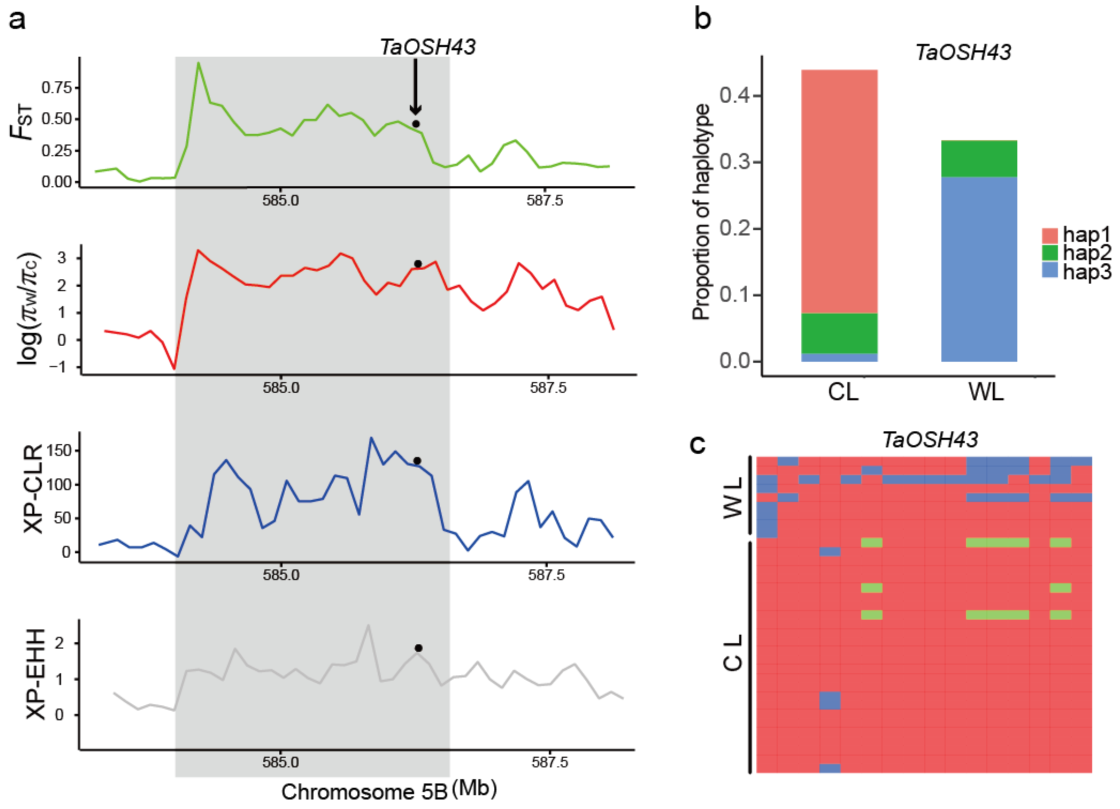


**Supplementary Figure 13**

**Population sweep selection between the Chinese landraces and West Asian landraces during their local adaption.**

(**a**) One gene distributed in a candidate region were commonly identified using the $F_{ST}$-$\theta\pi$ ratio, XP-CLR, and XP-EHH methods. The black points indicate the location of the candidate gene, which had a strong selection signal. (**b**) The haplotype frequency of the candidate gene between Chinese landraces (CL) and West Asian landraces (WL). (**c**) Heatmap of the Chinese and West Asian landrace genotypes in the putative selective sweep containing *TaOSH43*.

### Supplementary Tables 1 - 12

**Supplementary Table 1.** Wheat landraces and cultivars used in this study and their origins.

**Supplementary Table 2.** Summary of sequencing results.

**Supplementary Table 3.** Summary of the SNPs located in high-confidence genes.

**Supplementary Table 4.** The molecular diversity of the wheat population.

**Supplementary Table 5.** SNP heterozygosity of the wheat accessions.

**Supplementary Table 6.** Correlation between MAF and Tajima's *D*.

**Supplementary Table 7.** Candidate genes in the introgression regions. (**a**) Statistics of similar regions between the three populations. (**b**) Candidate genes introgressed from the European varieties to the Chinese cultivars within the candidate region located on chromosome 5D. (**c**) Candidate genes introgressed from the Chinese landrace to the Chinese cultivars within the candidate region located on chromosome 6A. (**d**) KEGG analysis of the introgressed genes from the European varieties to the Chinese cultivars in the candidate region located on chromosome 5D. (**e**) KEGG analysis of the introgressed genes from Chinese landraces to Chinese cultivars in the candidate region located on chromosome 6A.

**Supplementary Table 8.** Candidate genes from cultivars compared with landraces. (**a**) Genomic regions selected in the cultivar varieties. (**b** to **d**) Candidate selection regions in cultivars determined using the (**b**) $F_{ST}$-$\theta\pi$ method, (**c**) XP-CLR method, or (**d**) XP-EHH method. (**e**) Candidate selected genes in the cultivars commonly identified using the $F_{ST}$-$\theta\pi$, XP-CLR, and XP-EHH methods. (**f**) Candidate selected genes in the cultivars identified using only one of the $F_{ST}$-$\theta\pi$, XP-CLR, or XP-EHH methods. (**g**) KEGG analysis of the selected genes in the cultivars. (**h**) GO analysis of the selected genes in the cultivars. (**i**) Duplicated copies of the homoeologous genes selected in the cultivars.

**Supplementary Table 9.** Candidate genes from the Chinese cultivars compared with European landraces. (**a**) Genomic regions selected in the Chinese cultivars. (**b** to **d**) Candidate selection regions in the Chinese cultivars determined using the (**b**) $F_{ST}$-$\theta\pi$

method, (**c**) XP-CLR method, or (**d**) XP-EHH method. (**e**) Candidate selected genes in the Chinese cultivars commonly identified using the $F_{ST}$-$\theta\pi$, XP-CLR, and XP-EHH methods. (**f**) Candidate selected genes in the Chinese cultivars identified using only one of the $F_{ST}$-$\theta\pi$, XP-CLR, or XP-EHH methods. (**g**) KEGG analysis of the selected genes in the Chinese cultivars. (**h**) GO analysis of the selected genes in the Chinese cultivars. (**i**) Duplicated copies of the homoeologous genes selected in the Chinese cultivars.

**Supplementary Table 10.** Candidate genes from the European landraces compared with the West Asian landraces. (**a**) Genomic regions selected in the European landraces. (**b** to **d**) Candidate selected regions in the European landraces determined using the (**b**) $F_{ST}$-$\theta\pi$ method, (**c**) XP-CLR method, or (**d**) XP-EHH method. (**e**) Candidate selected genes in the European landraces commonly identified using the $F_{ST}$-$\theta\pi$, XP-CLR, and XP-EHH methods. (**f**) Candidate selected genes in the European varieties identified using only one of the $F_{ST}$-$\theta\pi$, XP-CLR, or XP-EHH methods. (**g**) KEGG analysis of the selected genes in the European landraces. (**h**) GO analysis of the selected genes in the European landraces.

**Supplementary Table 11.** Candidate genes from the Chinese landraces compared with the West Asian landraces. (**a**) Genomic regions selected in the Chinese landraces. (**b** to **d**) Candidate selected regions in the Chinese landraces determined using the (**b**) $F_{ST}$-$\theta\pi$ method, (**c**) XP-CLR method, or (**d**) XP-EHH method. (**e**) Candidate selected genes in the Chinese landraces commonly identified using the $F_{ST}$-$\theta\pi$, XP-CLR, and XP-EHH methods. (**f**) Candidate selected genes in the Chinese landraces identified using only one of the $F_{ST}$-$\theta\pi$, XP-CLR, or XP-EHH methods. (**g**) KEGG analysis of the selected genes in the Chinese landraces. (**h**) GO analysis of the selected genes in the Chinese landraces.

**Supplementary Table 12.** Primers for PCR. (**a**) Primer sequences. (**b**) The haplotypes of 28 Chinese cultivars determined using PCR.
